# Neuropilin-1 facilitates SARS-CoV-2 cell entry and provides a possible pathway into the central nervous system

**DOI:** 10.1101/2020.06.07.137802

**Authors:** Ludovico Cantuti-Castelvetri, Ravi Ojha, Liliana D. Pedro, Minou Djannatian, Jonas Franz, Suvi Kuivanen, Katri Kallio, Tuğberk Kaya, Maria Anastasina, Teemu Smura, Lev Levanov, Leonora Szirovicza, Allan Tobi, Hannimari Kallio-Kokko, Pamela Österlund, Merja Joensuu, Frédéric A. Meunier, Sarah Butcher, Martin Sebastian Winkler, Brit Mollenhauer, Ari Helenius, Ozgun Gokce, Tambet Teesalu, Jussi Hepojoki, Olli Vapalahti, Christine Stadelmann, Giuseppe Balistreri, Mikael Simons

## Abstract

The causative agent of the current pandemic and coronavirus disease 2019 (COVID-19) is the severe acute respiratory syndrome coronavirus 2 (SARS-CoV-2)^1^. Understanding how SARS-CoV-2 enters and spreads within human organs is crucial for developing strategies to prevent viral dissemination. For many viruses, tissue tropism is determined by the availability of virus receptors on the surface of host cells^2^. Both SARS-CoV and SARS-CoV-2 use angiotensin-converting enzyme 2 (ACE2) as a host receptor, yet, their tropisms differ^3-5^. Here, we found that the cellular receptor neuropilin-1 (NRP1), known to bind furin-cleaved substrates, significantly potentiates SARS-CoV-2 infectivity, which was inhibited by a monoclonal blocking antibody against the extracellular b1b2 domain of NRP1. NRP1 is abundantly expressed in the respiratory and olfactory epithelium, with highest expression in endothelial cells and in the epithelial cells facing the nasal cavity. Neuropathological analysis of human COVID-19 autopsies revealed SARS-CoV-2 infected NRP1-positive cells in the olfactory epithelium and bulb. In the olfactory bulb infection was detected particularly within NRP1-positive endothelial cells of small capillaries and medium-sized vessels. Studies in mice demonstrated, after intranasal application, NRP1-mediated transport of virus-sized particles into the central nervous system. Thus, NRP1 could explain the enhanced tropism and spreading of SARS-CoV-2.

## MAIN TEXT

An outbreak of SARS-CoV-2 infections started in December 2019 in the Chinese province of Hubei, causing a pandemic associated with a severe acute pulmonary disease named COVID-19 (coronavirus induced disease 2019)^1^. A related coronavirus, SARS-CoV, led to a much smaller outbreak in 2003, possibly because infection occurs predominantly in the lower respiratory system^6^. SARS-CoV-2, in contrast, spreads rapidly through active pharyngeal viral shedding^5^. Yet, uptake of both viruses is mediated by the identical cellular receptor, angiotensin-converting enzyme 2 (ACE2), present at the surface of some cells, mainly in the lower respiratory epithelium and other organs such as the kidney and the gastrointestinal tract^3,4^. The reason for the extended tissue tropism of SARS-CoV-2 with primary replication in the throat is unknown. One attractive hypothesis to explain differences in pathogenicity and tropism is the presence of a polybasic furin-type cleavage site, RRAR^S, at the S1–S2 junction in the SARS-CoV-2 spike protein that is absent in SARS-CoV^7^. Similar sequences are found in spike proteins of many other pathogenic human viruses, including Ebola, HIV-1 and highly virulent strains of avian influenza^7,8^. Insertion of a polybasic cleavage site results in enhanced pathogenicity by priming of fusion activity^9^ and create potentially additional cell surface receptor binding sites. In fact, proteolytic cleavage by furin, exposes a conserved carboxyterminal (C-terminal) motif RXXR_OH_ (where R is arginine and X is any amino acid; R can be substituted by lysine, K) on the substrate protein. Such C-terminal sequences that conforms to the ‘C-end rule’ (CendR) are known to bind to and activate neuropilin receptors (NRP1 and NRP2) at the cell surface^10,11^. Physiological ligands of NRPs, such as vascular endothelial growth factor A (VEGFA), and class-3 semaphorins (SEMA-3), are cleaved by furin and proprotein convertases, with the resulting C-terminal exposure triggering the receptor interaction^12^. These ligands are responsible for angiogenesis, and for mediating neuronal axon guidance, respectively^12^. Importantly, recent cryo-electron microscopy structure of the SARS-CoV-2 spike demonstrated the S1/S2 junction in a solvent-exposed loop, therefore accessible for interactions^13,14^.

To determine whether SARS-CoV-2 uses NRP1 for virus entry, we generated replication-deficient lentiviruses pseudotyped with SARS-CoV-2 spike protein (S) that drive expression of green fluorescent protein (GFP) upon infection. Such SARS-CoV-2 pseudoviruses are ideally suited for virus entry assays, as they allow separating viral entry from other steps in the virus life cycle, such as replication and assembly. Lentiviruses pseudotyped with the vesicular stomatitis virus (VSV) spike G were used as controls. For this assay, we used HEK-293T cells, as they do not express ACE2 and were not infected by lentiviral particles pseudotyped with the SARS-CoV-2 S protein. To test the potential role of NRP1 in virus infection, cells were transfected with plasmids encoding the main attachment receptors, either ACE2 or the transmembrane protease serine 2 (TMPRSS2), and NRP1. Both ACE2 and TMPRSS2 are required for cleavage of the S protein, and thus necessary for the fusion of viral and cellular membranes^9^. When expressed alone, ACE2 rendered cells susceptible to infection (Fig. 1a). NRP1 alone allowed lower, yet detectable levels of infection, both in HEK-293T and in Caco-2 cells (Fig. 1a,b), while cells transfected with plasmids encoding only TMPRSS2 were not infected (Fig. 1a). The co-expression of TMPRSS2 with either ACE2 or NRP1 potentiated the infection, with ACE2 together with TMPRSS2 being twice as efficient as NRP1 with TMPRSS2 (Fig. 1c). Maximal levels of infection were achieved when all three plasmids, driving expression of ACE2, NRP1 and TMPRSS2, were used for co-transfection (Fig. 1d). To further test the specificity of NRP1-dependent virus entry, we developed a series of monoclonal antibodies (mAbs), including function-blocking antibodies, against the extracellular b1b2 domain of NRP1, known to mediate the binding of CendR peptides^12^. The potency of these mAbs in preventing cellular binding and internalization of NRP ligands was tested using 80 nm silver nanoparticles (AgNP) decorated with the prototypic NRP1-binding CendR peptide RPARPAR _OH_^10^. These synthetic AgNP-CendR, but not the control particles without peptides, bind to and are internalized efficiently into PPC1 cells, which express high levels of NRP1^10^. One of the three antibodies, mAb3, efficiently blocked AgNP-CendR binding (Extended Data Figure 1a) and internalization (Extended Data Fig. 1b), while mAb1 had no effect and was used as a control in further experiments. mAb3 antibody recognizes the CendR binding pocket on the b1 domain of NRP1, and its binding to NRP1 with mutated binding pocket is compromised. Similar results were obtained in NRP1-expressing HEK-293T cells (Extended data Figure 2a,b). Importantly, upon incubation with HEK-293T cells expressing ACE2, NRP1 and TMPRSS2, mAb3 significantly reduced infection by SARS-CoV-2 pseudoviruses (Fig. 1e). To provide supporting evidence for the direct binding of virus particles to NRP1, SARS-CoV-2 pseudoviruses were pre-incubated with recombinant, soluble extracellular b1b2 domain of NRP1. The rationale of this approach was that the soluble form of NRP1 b1b2 would bind the virus particles and compete for the binding of NRP1 at the cell surface. As a negative control, we used b1b2 mutant with triple mutation (S346A, E348A and T349A in the CendR binding pocket), known to abrogate the binding of CendR peptides. We validated the effect of the recombinant NRP1 proteins, by showing that b1b2, but not the mutant protein, blocked uptake of AgNP-CendR in HEK-293T cells (Extended Data Fig. 2c). Importantly, addition of the soluble b1b2 domain of NRP-1 significantly reduced SARS-CoV-2 pseudovirus infection, whereas triple b1b2 mutant had no effect (Fig. 1f). Thus, these data demonstrate that NRP1 specifically potentiates SARS-CoV-2 pseudovirus infection.

**Figure 1.**
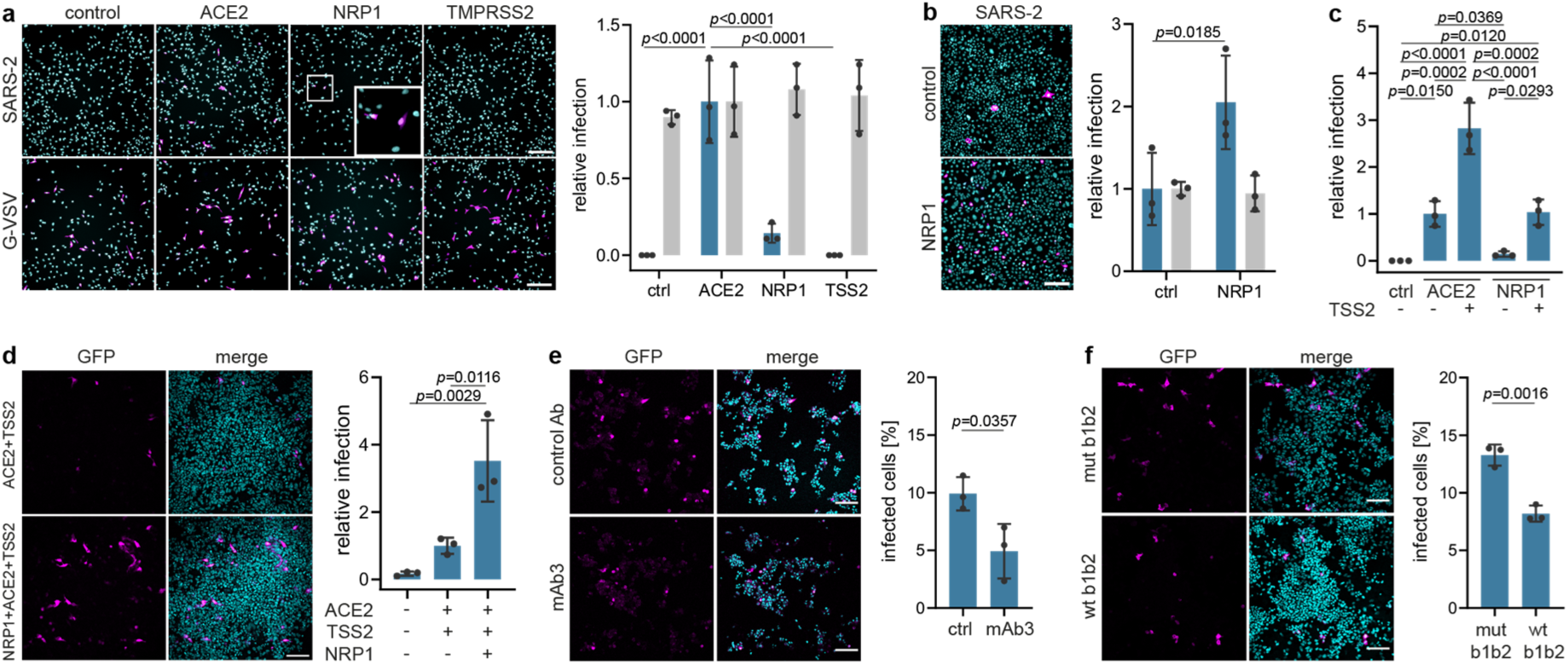
NRP1 facilitates the cellular entry of SARS-CoV-2 pseudotyped particles. **a**, Representative images and quantification of SARS-CoV-2 spike protein (SARS-2) (blue bars) and VSV-G pseudotype (grey bars) infectivity in HEK-293T cells expressing empty vector (control, ctrl), ACE2, NRP1 or TMPRSS2 (TSS2). Data are normalized to the respective infectivity of ACE2-expressing cells. Two-way ANOVA with Tukey’s correction for multiple comparisons. **b**, Representative image of SARS-2 infectivity of NRP1-expressing and empty vector-expressing Caco-2 cells. Quantification of SARS-2 and VSV-G pseudotypes infectivity in these cells. Data are normalized to control cells. Two-way ANOVA with Sidak’s correction for multiple comparisons. **c**, Infectivity of the SARS-2 pseudotype in ACE2 or NRP1-expressing HEK-293T cells, with and without TMPRSS2. One-way ANOVA with Tukey’s correction for multiple comparisons. **d**, Representative images and quantification of HEK-293T cells expressing ACE2+TMPRSS2 (upper panel) and NRP1+ACE2+TMPRSS2 (lower panel) after SARS-2 pseudotype inoculation. Data are normalized to ACE2+TMPRSS2-expressing cells. One-way ANOVA with Tukey’s correction for multiple comparisons. **e**,**f**, Representative images and quantification of HEK-293T cells expressing NRP1+ACE2+TMPRSS2 after SARS-2 pseudotype inoculation in the presence of mAb3 antibody against NRP1 (**e**, mAb3, lower panel) or control immunoglobulin (**e**, ctrl Ab, upper panel), and in the presence of NRP1 b1b2 domain (**f**, wt b1b2, lower panel) or the NRP1 mutant b1b2 domain (**f**, mut b1b2, upper panel). Two-tailed unpaired Student’s t test. All images show GFP-positive, infected cells (magenta) and Hoechst (cyan). Scale bars, 100 µm. All data is represented as mean ± s.d. from three independent experiments (a,b) or three biological replicates (d,e,f).

Next, we explored SARS-CoV-2 isolated from COVID-19 patients from the Helsinki University Hospital. RNA viruses such as SARS-CoV-2 have a remarkable ability to adapt to their host environment by generating adaptive mutations in a short period. Confirming recent reports^15^, we found that SARS-CoV-2 viruses, passaged in VeroE6 cells rapidly accumulated mutations around the furin cleavage site of the S protein that impaired its furin cleavage (Fig. 2a,b). We compared the effect of the blocking antibodies on infection of Caco-2 cells with wild-type and mutated SARS-CoV-2 virus, and found that Caco-2 cells pre-incubated with NRP1 blocking antibody reduced infection with wild-type virus by ∼40%, while the control antibody had no effect (Fig. 2c,d). In contrast, NRP1 blocking antibodies did not reduce the infection of Caco-2 cells with the mutated virus.

**Figure 2.**
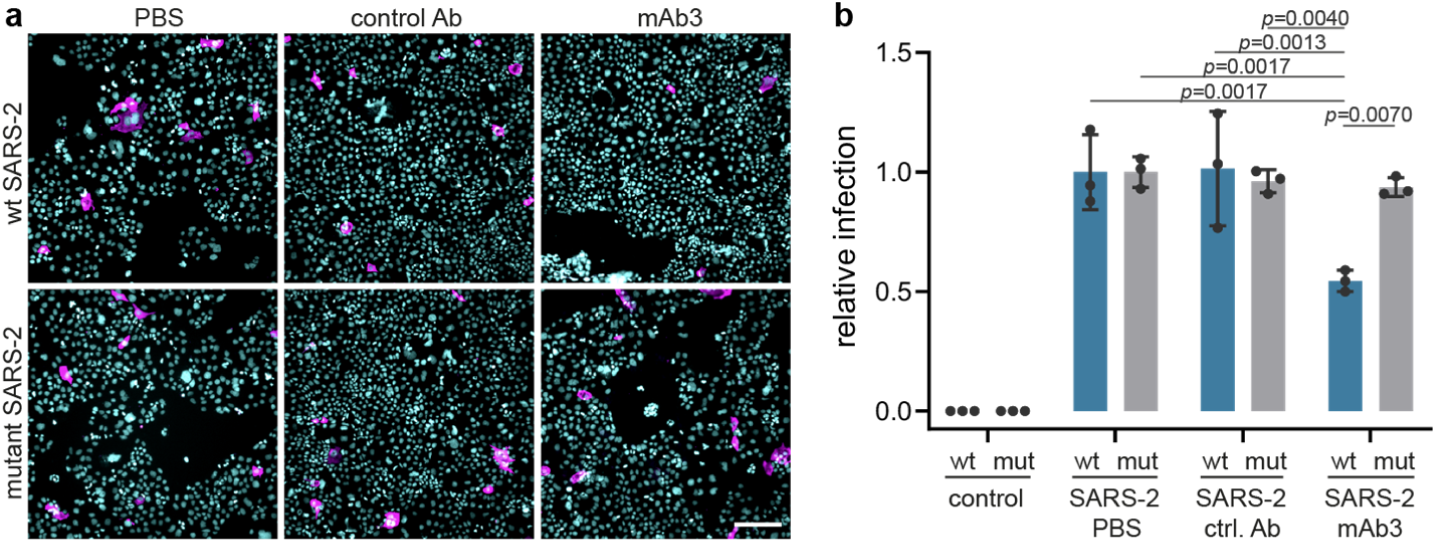
A blocking antibody against the b1b2 domain of NRP1 reduces infection of wild-type SARS-CoV-2 (wt SARS-2), but not a mutant with a deletion at the furin-cleavage site (mut SARS-2). **a**, Sequence analysis of viruses isolated at different passages (P) from different cell types. The first sequence is the reference from the Wuhan isolate (NC_045512.2). The sequence abundance in each virus population is indicated as a percentage (%). b, Deletion around the furin-cleavage site abrogates the furin cleavage of spike protein (S). Western blot analysis of cell lysates from VeroE6 cells infected for 16h with two viral populations (§ and §§). In lysates from cells infected with wt SARS-2, two bands are visible, the full-length S (upper band) and S1 (lower band). In lysates from cells infected with the mut SARS-2, only the uncleaved S is detected. **c**,**d**, Representative images (**c**) and quantification (**d**) of infection assays in Caco2 cells in the presence of control mAb1 (ctrl. Ab) or mAb3 blocking antibodies against NRP1 after wt SARS-2 (wt, blue bars) or mutant SARS-2 (mut, grey bars) inoculation Data are normalized to the respective vehicle (PBS) control sample. Mean ± s.d. from three independent experiments. SARS-2 and mutant SARS-2 infected cells (magenta) and Hoechst (cyan). Scale bar, 50 µm

Having obtained evidence that NRP1 facilitates SARS-CoV-2 entry, we examined whether NRPs are expressed in infected cells. For these analyses, we used published scRNA-seq datasets of cultured experimentally infected human bronchial epithelial cell (HBECs) and cells isolated from bronchoalveolar lavage fluid (BALF) of severely affected COVID-19 patients^16,17^. Using these datasets, we analyzed which of the proposed SARS-CoV-2 cell entry and amplification factors were correlated with the detection of virus RNA in single cell transcriptomes. Of all the proposed factors only three, *NRP1, FURIN* and *TMPRSS11A*, were enriched in SARS-CoV-2 infected compared to non-infected cells (Extended Data Fig. 3). In addition, RNA expression of *NRP1* and *NRP2* was elevated in SARS-CoV-2-positive compared to bystander cells isolated from bronchoalveolar lavage fluids of severely affected COVID-19 patients (Extended Data Fig. 4). Severe COVID-19 disproportionately affects patients with diabetes^18^. We analyzed a cryopreserved human diabetic kidney single-nucleus RNA sequencing dataset^19^, and found that among 14 proposed SARS-CoV-2 cell-entry and amplification factors, only *NRP1* was significantly upregulated (Extended Data Fig. 5). Next, we analyzed the expression pattern of NRP1 by examining the Human Protein Atlas (https://www.proteinatlas.org)^20^, which revealed particularly high levels of *NRP1* expression in the epithelial surface layer, outlining the respiratory and gastrointestinal tracts (Extended Data Fig. 6). To further resolve expression of SARS-CoV-2 cell-entry receptors, we took advantage of published scRNA-seq datasets of human lung tissue^21^ and human olfactory epithelium^22^. In the lung, both *NRP1* and *NRP2* were abundantly expressed with highest expression in endothelial cells and detectable levels in all pulmonary cells (Extended Data Fig. 7). In the adult human olfactory epithelium, we observed abundant expression of both *NRP1* and *NRP2* in almost all detected cell types of the olfactory epithelium, in sharp contrast to *ACE2*, which was sparsely expressed in these datasets (Extended Data Fig. 8).

In light of the widely reported disturbance of sense of olfaction in a large fraction of COVID-19 patients^23^, and the enrichment of NRPs in the olfactory epithelium, we studied whether SARS-CoV-2 could infect the NRP1-positive cells in the olfactory epithelium in humans. Thus, we analyzed a series of autopsies from six COVID-19 patients and seven non-infected control autopsies for the presence of SARS-CoV-2 infection in the olfactory system (Fig. 3, Extended Data Table 1). Using antibodies against the spike protein, we detected infection in the olfactory epithelium of five out of six COVID-19 patients. Positive signal was particularly well visible in the cells bordering the nasal cavity, whereas no S antigen signal was detectable in the control autopsies (Fig. 3). The infected olfactory epithelial cells showed high expression of NRP1 (Fig. 3a). In the underlying tissue, we detected spike protein in NRP1-positive endothelial cells. Additional co-staining of the transcription factor, OLIG2, and spike protein indicated infection of late olfactory neuronal progenitors and/or newly differentiated olfactory neurons (Fig. 3b). Strikingly, within the brain, the olfactory bulb and tracts displayed immunoreactivity for the spike protein especially within NRP1-positive endothelial cells in small capillaries and medium-sized vessels (Fig. 3c).

**Figure 3.**
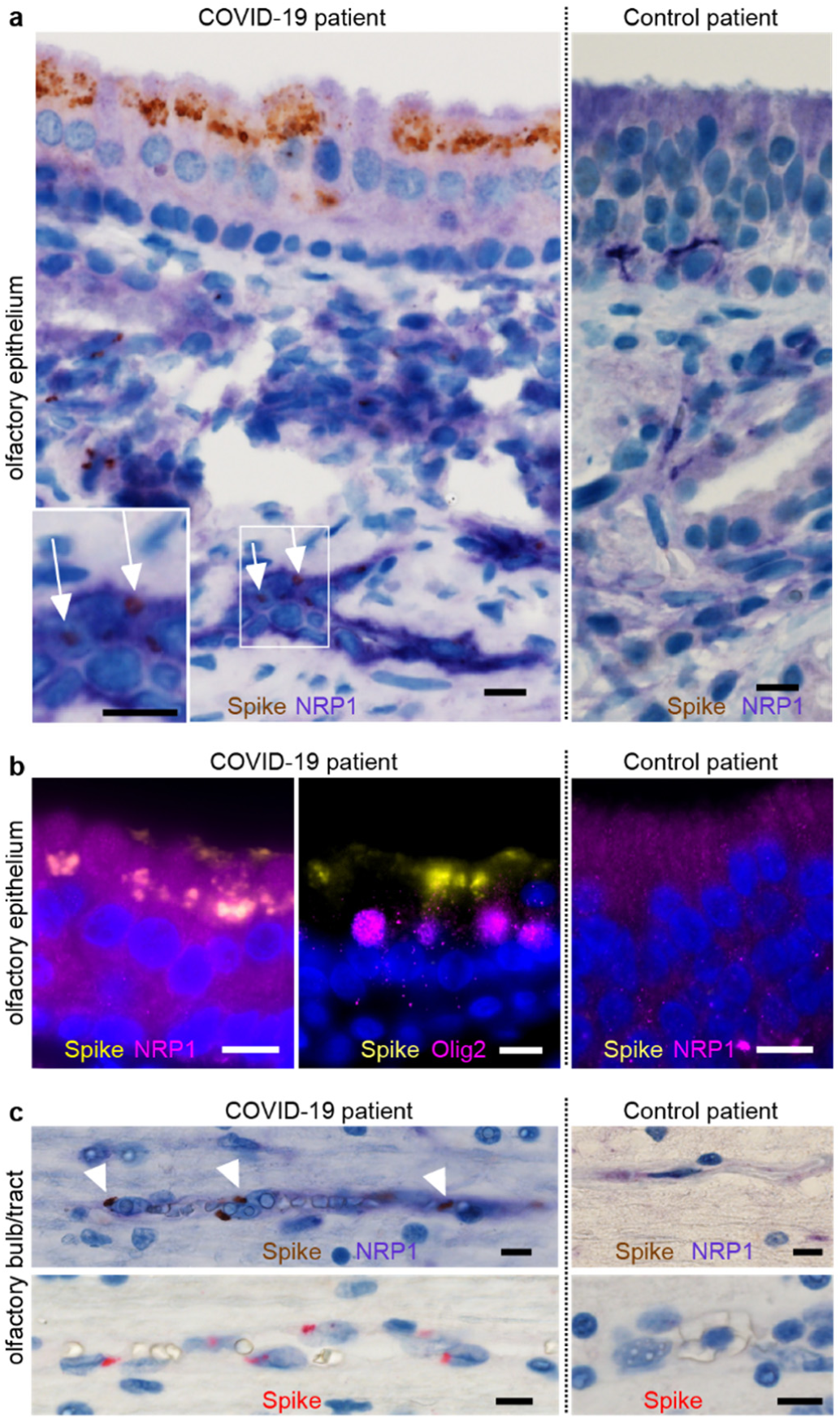
SARS-CoV-2 infects the olfactory epithelium and bulb. **a**, COVID-19 patient with abundant spike (S) protein immunoreactivity (IR) in the apical olfactory epithelium (OE) (brown) compared to non-infected control. Expression of NRP1 in OE cells and endothelia (liliac). Punctae of S protein (brown) indicate infection of endothelial cells (inset). **b**, Co-localization of NRP1 and S protein in OE cells (left, right). Co-staining of OLIG-2 (magenta) and S protein (yellow) reveals infection of late olfactory neuronal progenitors/newly differentiated olfactory neurons (middle). **c**, COVID-19 patient with S protein positive granules (brown) in NRP1 positive endothelial cells (liliac) in the olfactory bulb/tract compared to non-infected control (upper panel) and S protein IR alone (red, lower panel). Scale bar, 10 µm.

These results provide evidence that SARS-CoV-2 infect brain tissue, consistent with its multi-organ involvement^24^, and suggest that viral entry into the brain may occur through the olfactory epithelium. To determine whether NRPs could provide a pathway for virus entry into the brain through the olfactory system, we performed experiments in mice. First, we confirmed the high expression of NRPs in the mouse olfactory epithelium. Virtually all sensory olfactory neurons expressed NRP1 (Extended Data Fig. 9). To study whether NRPs could provide a transport pathway into the nervous system, we used the 80 nm diameter AgNPs-CendR, which were validated for their specific interaction with NRPs (Extended Data Fig. 2). AgNPs-CendR and control AgNPs lacking the peptide, both of similar size as many viruses, were administered into the nose of anesthetized adult mice (Fig. 4). Mice were sacrificed 30 min and six hours after intranasal administration and analysis of the olfactory epithelia revealed much larger uptake of AgNP-CendR particles compared to control particles into the epithelia (Fig. 4a-c). Next, we studied whether AgNP-CendR could be transported to the central nervous system. Strikingly, AgNP-CendR, but not control particles, were readily detected in brain tissue (Fig. 4a,c). Whereas AgNP-CendR were found particularly enriched in neurons located in the cortex, they were also found in the olfactory bulb, mainly in neuronal cells (Fig. 4b), and, to a lesser extent, in endothelial cells (Extended Data Fig. 10). These data provide evidence for the existence of a NRP-dependent intranasal brain entry pathway.

**Figure 4.**
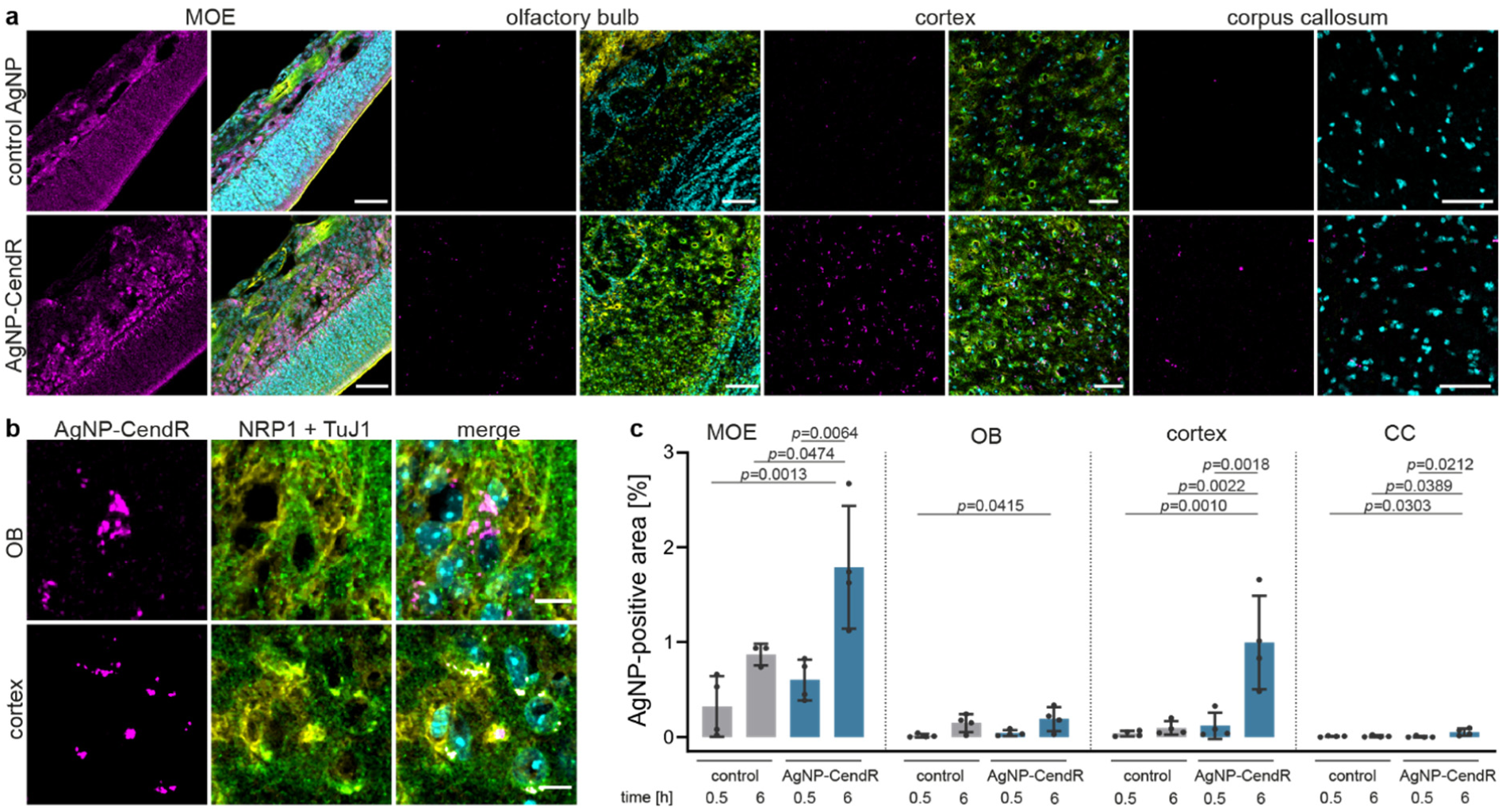
NRP1-dependent delivery of particles to the central nervous system. **a**, Representative images of main olfactory epithelium (MOE), olfactory bulb (OB), cortex and corpus callosum (CC) of animals treated for 6 hours with CF647-labeled silver nanoparticles (AgNP) conjugated with biotin (control AgNP, top row) or biotin-X-RPARPAR (AgNP-CendR, bottom row), showing the delivery and accumulation of particles. A small accumulation of AgNP-CendR was observed in the corpus callosum. Scale bar, 50 µm. **b**, Confocal images of OB and cortex of AgNP-CendR treated mice, showing that several AgNP-CendR^+^ cells (magenta) were positive for NRP1 (green) and TuJ1 (yellow). Scale bars, 10 µm. **c**, Quantification of the area of tissue occupied by AgNPs, showing a significant accumulation of AgNP-CendR in MOE, cortex and CC. *n* = 4 mice. Pseudocolored images, AgNP (magenta), NRP1 (green), TuJ1 (yellow), Hoechst (cyan). Data are means ± s.d. One-way ANOVA with Tukey’s correction for multiple comparisons within each group.

There is limited knowledge about the virus-host interactions that determine cellular entry and tissue spreading of SARS-CoV-2. So far, the focus has been entirely on ACE2 as a receptor, but it is clear that viruses display considerable redundancy and flexibility in receptor usage, in particular, because viruses can exploit weak multivalent interactions to enhance affinity^2^. Here, we provide evidence that NRP1 can serve as an entry factor, but more work is required to understand whether the engagement with NRP1 provides attachment, enhances proteolytic cleavage of S, induces receptor-mediated endocytosis, or promotes signaling. The reason why a number of viruses, such as the human T-cell lymphotropic virus type 1 (HTLV-1)^25^, cytomegalovirus (CMV)^26^, Epstein-Barr virus (EBV)^27^ and Lujo virus (LUJV)^28^, use NRPs as an entry factor could be due to its localization on epithelia facing the external environment, and enabling cell, vascular and tissue penetration^10^. The functions of NRPs are complex, and it is conceivable that some viruses do not use NRPs as bona fide endocytic receptors, but rather as signaling platforms to facilitate their spread. NRP1 ligation can increase vascular permeability and induce nutrient-sensitive macropinocytosis^11,29^. In this context, vascular endothelialitis, thrombosis, and angiogenesis that has been observed in COVID-19 patients, together with the upregulation of NRPs in SARS-CoV-2 infected blood vessels, is noteworthy^30^. Mechanistic details of how NRPs promote tissue penetration is under investigation, but the rapid infiltration of CendR peptides across tumor tissue implies active transport from cell to cell^10^. Such transcytotic pathways could be relevant for the transport of viruses from the olfactory sensory nerve endings along the axons into the brain, but paracellular pathways induced by enhanced vascular permeability are also conceivable. Proteolytic cleavage of viral spike proteins exposing CendR sequences are common to many neurotropic viruses (e.g. CMV, HTLV-2, measles, tick-born encephalitis, among others); thus, clarifying whether NRPs are involved in the neuro-invasive potential of furin-activated viruses is an important topic for future research.

## METHODS

### Animals

All animal studies were performed in compliance with the animal policies of the German Center for Neurodegenerative Diseases (Munich), and were approved by the District Government of Upper Bavaria. Animals were group-housed (3-5 mice) with 12 hour dark/light cycle and had access to food and water *ad libitum*. Adult male and female C57BL/6N mice (8 to 10 weeks of age) were taken for all experiments. Male mice were subjected to intranasal delivery of the particles. BALB/c mice were purchased from Jackson Laboratories.

### Plasmids

pQCXIBL-hTMPRSS2 and pCG1-SARS-2-S-HA were a kind gift from S. Pöhlmann (German Primate Center, Göttingen, Germany)^31^. hACE2 was a gift from H. Choe (Addgene plasmid # 1786). pCG1-MCS, pCG1-ACE2, pCG1-NRP1 were cloned by PCR amplification of vector and insert, followed by 2-fragment Gibson assemblies. PCRs were performed using Q5 polymerase (New England Biolabs Inc) according to the manufacturer’s protocol. Fragments were generated with the following primers (template plasmids indicated in brackets): pCG1-MCS: 5’ fragment (pCG1-SARS-2-S-HA): 5’-gctcagtggaacgaaa-3’ (fwd) and 5’-aacagtcgactctagaggatccgaattcgcc-3’ (rev), 3’fragment: (pCG1-SARS-2-S-HA): 5’-gcgaattcggatcctctagagtcgactgtttaaac-3’ (fwd) and 5’-ccttaacgtgagttttcgt -3’(rev), pCG1-ACE2: vector (pCG1-SARS-2-S-HA): 5’-gtcgactgtttaaacctgc-3’ (fwd) and 5’-aacgctagccagcttccgaattcgccctatag-3’ (rev), insert (hACE2): 5’-atagggcgaattcggaagctggctagcgt-3’ (fwd) and 5’-cttgcatgcctgcaggtttaaacagtcgacggatccaagcttaagcg-3’ (rev), pCG1-NRP1: vector pCG1-SARS-2-S-HA): 5’-agagtcgacctgcag-3’ (fwd) and 5’-gatcgctccttactaccgaattcgccctat-3’ (rev), insert (NRP1, native human ORF in pF1K, FXC23812, Promega): 5’-atagggcgaattcggtagtaaggagcgatcgc-3’ (fwd) and 5’-gatcagcttgcatgcc-3’ (rev). Gibson assembly was performed using NEBuilder® HiFi DNA Assembly Cloning Kit (New England Biolabs Inc.) according to the manufacturer’s recommendations. All constructs were verified by Sanger sequencing. The experiments described in Figure 1a, b and c, and in Figure 2 were performed using the following plasmids: pLenti-6.3-hNRP1, pLenti-6.3-hACE2, pLenti-6.3 hTMPRSS2 and pLenti-6.3 (vector) obtained by the Genome Biology Unit core facility of the University of Helsinki. The plasmids where generated by Gateway cloning from a genome-wide orfeome library available in house (https://www.helsinki.fi/en/researchgroups/genome-biology-unit/clones-and-cloning). Genome Biology Unit is supported by HiLIFE and the Faculty of Medicine, University of Helsinki, and Biocenter Finland. All plasmids were sequenced using standard CMV-forward and V5-reverse primers.

### Cell culture

HEK 293T and Caco-2 (ATCC) cells were grown in complete growth media supplemented with 10% fetal calf serum (FCS), (pen/strep, L-Glutamine in DMEM) and passaged 1:8 (HEK-293 T) or 1:5 (Caco-2) every three days. PPC-1 human primary prostate cells were obtained from Erkki Ruoslahti’s laboratory at the Cancer Research Center, Sanford-Burnham-Prebys Medical Discovery Institute. Cells were cultivated as an attached culture in high-glucose Dulbecco’s Modified Eagle Medium (DMEM; Lonza) containing 100 IU/ml of streptomycin and penicillin (both Thermo Fischer Scientific), and 10% fetal bovine serum (FBS; Thermo Scientific) in a 37°C incubator with 5% CO2. For the transfection of NRP1, ACE2 and TMPRSS1, 8 mm glass coverslips were coated with 0,01% poly-L-lysine for 1 hour at 37°C and 5×10^5^ cells were plated in each well of 24 well plates. Cells were also plated on 96-well imaging plates (PerkinElmer) at a dilution of 1×10^4^ cells per well in 100ul of full growth medium. 24 hours after plating, cells were transfected in Opti-MEM supplemented with GlutaMAX (Gibco) and 5% FCS with Lipofectamine 3000 reagent (Thermo Fisher Scientific) or Transit LT-1 (Mirus) according to the manufacturer’s instructions. Briefly, 0.25 µg (24 well plates) or 0-10 µg (96 well plates) of DNA per well was prepared in Opti-MEM and P3000 reagent (for Lipofectamine 3000) or Opti-MEM (for Transit LT-1). The DNA was then mixed with the transfection reagent and left at room temperature for 5 minutes (for Lipofectamine) or 15 minutes (for Transit LT-1). Cells were then incubated with the mix for 4 hours before the transfection reagent was replaced with fresh growth media. The viral particles were added 24 hours after transfection. After treatment, the cells were fixed in 4% paraformaldehyde (PFA) for 20 min. After washes with PBS, the cell nuclei were counterstained with Hoechst 33342 DNA stain solution (Thermo Fisher Scientific) and mounted with ProLong Gold Antifade (8 mm glass coverslips) or left in PBS (96-well plates) before imaging at SP5 confocal microscope or Molecular Device Image Xpress Nano high content imaging microscope, respectively. For confocal imaging of NRP-1 expressing PPC-1 cells, 30,000 cells/well were cultured at 37°C in 5% CO_2_ as a monolayer on coverslips (d = 12 mm; Paul Marienfeld GmbH & Co. KG) in 24 well plates (#08-772-1H, Thermo Fisher Scientific) with DMEM. The attached cells were pre-incubated at 37°C for 30 min with mAbs (150 μl, 30 μg/ml), which were dialyzed prior to use overnight against 1X PBS using 10 kDa dialysis cassettes (#66380, Thermo Fischer Scientific). Next, RPARPAR-targeted or non-targeted CF555-labeled AgNPs (1.5 nM final concentration) in supplemented DMEM were added to the wells, and the cells were further incubated at 37°C for 1 h. Next, the cells were washed 3 times with PBS, fixed with −20 °C methanol (MeOH; Naxo) for 5 min, and counterstained with DAPI (500 μl, 1 μg/ml; Thermo Fischer Scientific). Microscopy slides (76 × 26 mm, Glaswarenfabrik Karl Hecht GmbH & Co. KG) were coverslipped using Fluoromount-G aqueous mounting medium (Electron Microscopy Sciences). Confocal imaging was performed using an Olympus FV1200MPE confocal microscope (Olympus Europa SE & Co. KG) with a UPlanSApo 60x/1.35na objective (Olympus Europa SE & Co. KG), and images were analyzed with FluoView FV10-ASW 4.0 software (Olympus Europa SE & Co. KG).

### Production of SARS-CoV-2 S-pseudotyped lentiviral particles

HEK-293T cells were grown in complete growth media (10% FCS, Pen/strep, L-Glutamine in DMEM) until 60 to 70% confluent in a 10 cm dish. The medium was then replaced with Opti-MEM (supplemented with L-Glutamine and 5% FCS, no antibiotics) for two hours. Cells were then transfected with Lipofectamine 3000 according to the manufacturer’s instructions. Briefly, a solution with Lipofectamine 3000 and a solution with a mix of 18 µg of the following plasmids were separately prepared in Opti-MEM: pMDL g/p RRE, pRSV-REV, pLenti-GFP and a plasmid encoding the spike proteins of SARS-CoV-2 or VSV-G. Plasmid ratio for pMDL g/p RRE, pRSV-REV, the spike protein plasmid and pLenti-GFP is 1:1:1:2, respectively. The P3000 reagent was then added to the mix. After 5 minutes, the DNA mix was added to Lipofectamine solution and the mix was left for 20 minutes at room temperature. The mix was then added to the cells for 6 hours. Cell medium was then changed to normal growth medium (without antibiotics). 24 and 48 hours after transfection, the medium was collected, centrifuged at 1000g for 5 minutes, and filtered through a 0.45 µm filter. The medium was centrifuged at 65000g with a SW28 rotor for 3 hours over a cushion of 10% sucrose prepared in PBS. The pellet was finally resuspended in TBS buffer containing 5% bovine serum albumin overnight at 4°C, aliquoted and frozen at -80°C.

### Recombinant NRP1 b1b2 soluble protein expression and purification

Wild type and triple mutant human NRP1 b1b2 domain (residues 274-584) were expressed in *Escherichia coli* strain Rosetta-gami-2 (Novagen, Madison, WI) as a His-tag fusion in pET28b (Novagen). Cells were grown in Terrific-Broth at 37°C to an OD600 = 1.2 and after 15 min at 4°C induced with 1 mM isopropyl β-d-thiogalactoside. After growth at 16°C for 16 h, cells were harvested by centrifugation, lysed, and centrifuged, and proteins were purified over HIS-Select (Sigma–Aldrich, St. Louis, MO) nickel affinity resin in 20 mM Tris (pH 8.0) and 400 mM NaCl with an imidazole gradient from 25-500 mM. Further purification was performed on 5 ml hitrap heparin column (GE). Protein was loaded in 20mM Tris pH=8.0, 100mM NaCl and eluted using a linear gradient 100-800 M NaCl. Final purification was performed using Superdex 75 16/100 (Amersham Pharmacia) column equilibrated in 20mM Tris pH 8.0, 150mM NaCl.

### Generation of monoclonal antibodies against NRP1 b1b2

Female BALB/c and C57BL/6 mice, 8–9 weeks old, were immunized intraperitoneally with 17 μg of recombinant NRP1 b1b2 mixed with an equal volume of complete Freund’s adjuvant (Sigma–Aldrich Chemie, Steinheim, Germany), followed by a booster immunization four weeks later of the same dose mixed with incomplete Freund’s adjuvant (Sigma–Aldrich). Mice received three boosts of the same amount of antigen in PBS on days −3, −2, and −1 prior to fusion. Spleens were excised and the splenocytes were fused with myeloma cells (P3×63Ag8.653) according to a previously described protocol^32^. Beginning on day 10 after fusion, hybridoma supernatants were screened for specific antibodies. Before experiments, the hybridoma supernatants were centrifuged at 300g for 5 min at room temperature and 500 µl dialyzed against 2 L of PBS over night at 4°C prior use.

### Synthesis and Functionalization of AgNPs

The silver nanoparticles (AgNPs) were synthesized and functionalized as described previously ^33^, wherein biotin-Ahx-RPARPAR-OH (RPARPAR; TAG Copenhagen A/S) was used as the targeting moiety and CF555 dye (#92130, Biotium) or CF647 (#92135, Biotium) as the fluorophore. Briefly, AgNO_3_ (360 mg; #209139, Sigma-Aldrich) was added to ultrapure Milli-Q (MQ) water (2 L; resistivity 18 MΩ cm^−1^) in a flask cleaned with a piranha solution (H_2_SO_4_/H_2_O_2_). Next, trisodium citrate hydrate (400 mg; #25114, Sigma-Aldrich) was dissolved in MQ water (40 ml) and added to the vessel. The solution was boiled for 30 min in the dark. The resulting Ag-citrate was used directly in the next step. Next, NeutrAvidin (NA; #31055, Thermo Scientific) was modified with a OPSS-PEG(5K)-SCM linker (OPSS; JenKem Technology USA) according to the procedure described by Braun *et al*^33^. Subsequently, NeutrAvidin-OPSS (3.9 ml, 2.9 mg/ml) was added to Ag-citrate (500 ml). After 2 min, 4-morpholineethanesulfonic acid hemisodium salt (5 ml, 0.5 M in MQ water; #M0164, Sigma-Aldrich) was added. The pH of the solution was adjusted to 6.0, and the solution was kept at 37°C for 24 h. The solution was brought to room temperature (RT) and 10X phosphate buffered saline (50 ml; PBS; Naxo) was added, followed by Tween® 20 (250 µl; #P9416, Sigma-Aldrich). The solution was centrifuged at 12,200 × g for 1 h (4°C), the supernatant was removed, and the particles were resuspended in PBST (0.005% Tween® 20 in PBS). Next, tris(2-carboxyethyl)phosphine hydrochloride solution (TCEP; #646547, Sigma-Aldrich) was added to a final concentration of 1 mM, followed by a 30 min incubation at RT. Then lipoic acid-PEG(1k)-NH_2_ (#PG2-AMLA-1k, Nanocs) was added to a final concentration of 5 µM, and the mixture was incubated at RT for 2.5 h. The solution was centrifuged at 17,200 × g for 20 min (4°C), the supernatant was removed and the particles were resuspended in PBST to half of the initial volume. The AgNP solution was filtered through a 0.45 µm filter and stored at 4°C in dark. NHS-functionalized CF555 or CF647 dye was coupled to the NH_2_ groups of the linker on the AgNPs. For this, NHS-CF555 (5 µl, 2 mM) in dimethyl sulfoxide (DMSO; Sigma-Aldrich Co., LLC) was added to AgNP (500 µl), followed by an overnight incubation at 4°C. The particles were washed 3 times by centrifugation at 3,500 × g for 10 min at 4°C, followed by resuspension of the particles in PBST by sonication. Next, biotinylated RPARPAR peptide were coupled to the particles by adding peptide (10 µl, 2 mM in MQ water) to AgNPs (500 µl), followed by incubation at RT for 30 min. The AgNPs were washed, 0.2 µm filtered and stored at 4°C in dark.

### Mouse Immunohistochemistry and Immunofluorescence

The mouse olfactory epithelium, the olfactory bulbs, and the brain were isolated and left in 30% sucrose until sunk to the bottom of the tube. After embedding in Tissue Tek OCT, the tissues were sectioned with a thickness of 14 µm. For the multiplex fluorescence labeling of NRP1 mRNA (RNAScope Probe-Mm-Nrp1, Advanced Cell Diagnostics, 471621), the RNAscope^®^ Multiplex Fluorescent v2 Assay (Advanced Cell Diagnostics, 323100) was used according to the manufacturer’s instructions. For the immunohistochemistry, the sections were air dried overnight at room temperature, washed in PBS and permeabilized with 0.1% Triton-X 100 for 15 min at room temperature. The blocking of non-specific staining was performed with a blocking solution of 1% fetal calf serum, 1% fish gelatin and 1% bovine serum albumin in PBS for one hour at room temperature, followed by a second blocking step for endogenous mouse Ig with AffiniPure Fab fragment Donkey anti-mouse IgG (Biozol, JIM-715-007-003) diluted 1:100 in blocking solution for one hour at room temperature. Primary antibodies were diluted in 10% blocking solution: 1:250 NRP1 (monoclonal rabbit, ab81321, Abcam); 1:1000 TuJ1 (monoclonal mouse, G712A, Promega); 1:250 NeuN (polyclonal chicken, ABN91, Milipore); 1:2000 GFP (polyclonal rabbit, A-6455, Thermo Fisher Scientific), 1:250 ColIV (polyclonal goat, 1340-01, Southern Biotech), 1:250 CD31 (monoclonal mouse, sc-376764, Santa Cruz Biotechnology), and incubated overnight at 4°C. After three washes in PBS, the sections were incubated in secondary antibody: Alexa Fluor 488 donkey anti-mouse (R37114, Thermo Fischer Scientific); Alexa Fluor 488 goat anti-rabbit (A21428, Thermo Fischer Scientific); Alexa Fluor 488 goat anti-chicken (A11039, Thermo Fischer Scientific); Alexa Fluor 488 donkey anti-goat (A11055, Thermo Fischer Scientific); Alexa Flour 555 donkey anti-goat (A21432, Thermo Fischer Scientific); Alexa Flour 555 goat anti-mouse (A21422, Thermo Fischer Scientific); Alexa Flour 647 donkey anti-rabbit (A31573, Thermo Fischer Scientific), Alexa Fluor 488 Phalloidin (A12379 Thermo Fischer Scientific), for two hours at room temperature. After three washes in PBS, the sections were counterstained with Hoechst 33342 stain solution (Thermo Fisher Scientific) and mounted with Prolong Gold Antifade Mountant (Thermofisher, P36930). For immunofluorescence, the cells were permeabilized on ice for 10 minutes in a solution of 0.1% Triton-X 100 and 0.1% Sodium citrate in PBS, blocked for one hour in blocking solution at room temperature and left overnight at 4°C in primary antibody diluted in 10% blocking solution. After three washes in PBS, the cells were incubated with secondary antibodies (diluted 1:500 in 10% blocking solution) for one hour at room temperature. After washes in PBS, the cells were counterstained with Hoechst 33342 (Thermo Fisher Scientific) stain solution and mounted with ProLong Gold Antifade. The stained tissues and cells were imaged on a Leica SP5 confocal microscope.

### Human immunohistochemistry

All experiments with human materials were approved by the ethics committee of the University Medical Center Göttingen and were performed in accordance with the respective national, federal and institutional regulations. Human tissue blocks were fixed in 3.7% formalin overnight and embedded in paraffin. 3 µm deparaffinized tissue sections were pretreated for epitope retrieval as indicated. Sections were treated with 3% H2O2 and permeabilized with 1% Triton-X 100 and then blocked with PBS containing 10% goat serum. Primary antibodies were applied over night at a dilution of 1:100 for SARS-CoV S protein (monoclonal mouse, ab272420, Abcam; microwave, citric acid buffer, 10 mM, pH 6.0), 1:250 for NRP1 (monoclonal rabbit, ab81321, Abcam; microwave, Tris-EDTA, pH 8.0) and 1:150 for OLIG2 (polyclonal rabbit, 18953, IBL; microwave, Tris-EDTA, pH 8.0). Secondary antibodies were added as follows: biotinylated anti-mouse 1:200 (GE Healthcare RPN 1001) followed by Tyramide Super Boost with Alexa Fluor 488 1:500 (Thermo Fisher Scientific) and Alexa Fluor 555 anti rabbit 1:500 for 2 h, at room temperature. Coverslips were mounted with ProLong Diamond Antifade with DAPI (Thermo Fisher Scientific). Either the Envision+ System for mice (Dako) and alkaline phosphatase conjugated anti-rabbit antibodies with FastBlue (Sigma Aldrich) or alkaline phosphatase coupled anti-mouse antibodies with FastBlue (Sigma Aldrich) were used for immunohistochemistry. Images were taken using a fluorescence microscope (Olympus BX-63 with a colour camera (Olympus DP80). Image visualization was performed using Omero Server software 5.6, Omero figure 4.0.2 (https://github.com/ome/omero-figure) and InkScape 0.92^34^. The human NRP1 protein immunohistochemistry data on 12 tissue types corresponds to images on the HPA database http://www.proteinatlas.org^20^.

### Intranasal delivery of silver nanoparticles and pseudotyped particles in mice

Intranasal delivery was performed as previously described ^35^. Mice were anesthetised with a mix of metedomidine, midazolam and fentanyl (MMF) and positioned in a head down and forward position. 3 µl of the solution containing the viral particles (Tris buffer saline with 5% bovine serum albumine) or the silver nanoparticles was then positioned close to the nostril of the animals until inhaled. After 3 minutes, 3 µl of the solution containing the viral particles was delivered to the other nostril. This procedure was repeated until 30 µl of solution were delivered to each animal. The animals were then woken up with a mix of atipamezole, flumazenil, and naloxone (AFN). Analgesia was achieved with daily injection of buprenorphine. At the end of the experiment, the animals were anesthetised with ketamine and xylazine and perfused with PBS and 4% PFA.

### Image analysis

Images were analysed with CellProfiler 3.1.8 (https://cellprofiler.org/). Nuclei were identified automatically using the Otsu algorithm inbuilt in the software and the infected and non-infected cells were distinguished by thresholding the fluorescent signal of the SARS N protein immunostained cells. Non infected wells were used as negative controls. For each experiment in 96 plates, at least 2000 cells were analyzed. The numbers of NRP1^+^ positive neurons in the mouse MOE and brain were manually counted from three representative sections of three independent animals. All other measurements were performed semiquantitatively using Fiji software ^36^. To measure the infectivity of cultured cells, three randomly selected areas per coverslip were imaged. The percentage of GFP positive cells was determined by dividing the number of GFP^+^ nuclei by the number of Hoechst^+^ nuclei. The same threshold value was applied to all the coverslips of an independent experiment. To measure the AgNP uptake in culture cells, three randomly selected areas per coverslip were imaged. AgNP positive cells were determined the following way: each cell was identified by the positivity for the nuclear staining Hoechst 33342 and the actin-binding toxin phalloidin. Hoechst^+^ nuclei were used to define the cell ROI. Within each ROI, the area occupied by the signal of AgNPs was measured and expressed as area fraction. ROIs with an area fraction higher than 30% were considered AgNP^+^ cells. The percentage of AgNP^+^ cells was determined by dividing the number of AgNP^+^ cells by the number of Phalloidin^+^ cells. All samples were analyzed at the same threshold values. For the in vivo analysis three representative sections from four independent animals were stained with the specific antibodies. For each tissue a representative area was selected and the area occupied by AgNP particles with a size higher than 1.8 µm^2^ was measured. The same threshold value was applied to the images of each area.

### Isolation of WT SARS-CoV-2 from COVID-19 patient and virus propagation

Samples were obtained under the Helsinki University Hospital laboratory research permit HUS/32/2018 § 16. 500 µl of nasopharyngeal swab in Copan UTM® Universal Transport Medium was inoculated on Calu-3 cells and incubated for 1 h in +37°C, after which the inoculum was removed and replaced with Minimum Essential Medium supplemented with 2% FBS, L-glutamine, penicillin and streptomycin. Virus replication was determined by RT-PCR for SARS-CoV-2 RdRP ^37^, and the infectious virus titer was determined by plaque assay in VeroE6 cells. Viruses used in Fig. 2 were propagated in VeroE6 cells (SARS_mutFC_) or Caco-2 cells up to passage 7 or passage 1, respectively.

### Sequencing of virus isolates

Viral RNA was extracted from cell culture supernatants using QIAamp Viral RNA Mini Kit (Qiagen) and reverse-transcripted to cDNA using LunaScript RT SuperMix kit (New England Biolabs) following manufacturer’s instructions. Primer pools targeting SARS-CoV-2 were designed using PrimalScheme tool http://primal.zibraproject.org^38^ and PCR was conducted using PhusionFlash PCR master mix (ThemoFisher). Sequencing libraries were prepared using NEBNext ultra II FS DNA library kit (New England Biolabs) according to the manufacturer’s instructions, and sequenced using Illumina Miseq with v3 sequencing kit. Raw sequence reads were trimmed and low quality (quality score <30) and short (<25 nt) sequences removed using Trimmomatic^39^. The trimmed sequence reads were assembled to the reference sequence (NC_045512.2) using BWA-MEM algorithm implemented in SAMTools version 1.8^40^. The minority variants and insertion/deletion sites were called using LoFreq* version 2.1.4^41^.

### scRNA-seq analysis

Analyses of the scRNA-seq datasets including filtering, normalization and clustering were conducted using Seurat 3.1^42^. Human lung data from Han *et al*. ^21^, was downloaded from https://figshare.com/articles/HCL_DGE_Data/7235471, in the form of batch-corrected digital gene expression matrices and cell annotation csv files. Cell annotation included cell types, tissue of origin and age. Gene expressions were log-normalized with a scale factor of 10 000 using the NormalizeData function. Next, data was scaled using the ScaleData function and the number of UMI and the percentage of mitochondrial gene content were regressed out as described by the authors^21^. The first 40 principal components were considered for the UMAP (Uniform Manifold Approximation and Projection). Human olfactory neuroepithelium raw data from Durante *et al*.^22^, was downloaded from Gene Expression Omnibus under accession code GSE139522. Data was quality controlled, pre-processed, filtered, normalized, scaled and clustered as described in detail by the authors^22^. Data from 4 patients were integrated using the Seurat 3 standard integration^42^ using the parameters described by Durante *et al*.^22^. Since cell type annotations were not provided publicly, we have used the cell type specific markers indicated by the authors to identify cell type clusters. Differentially expressed genes table from Navindra *et al*., was obtained from the GitHub page maintained by the authors (https://github.com/vandijklab/HBEC_SARS-CoV-2_scRNA-seq). Table of differentially expressed genes (DEG) by cell type comparing the diabetic patients to the control patients, was obtained from Wilson *et al*.^19^. Bronchoalveolar lavage fluid (BALF) data was acquired from GSE145926 and only the scRNA-seq data was used ^17^. Datasets from 12 samples were quality controlled, filtered and integrated using the Seurat 3 standard integration workflow as described by the authors. Differentially expressed genes were identified comparing SARS-CoV-2 positive and negative cells with the FindMarkers function in Seurat. Cell annotation of infection status, cell type, etc. was provided by the authors on their GitHub page (https://github.com/zhangzlab/covid_balf). For all datasets except the DEG tables from Navindra *et al*.^16^ and Wilson *et al*.^19^, DotPlot and DimPlot functions within Seurat were used for visualization purposes.

### Statistics

All measurements were taken from distinct samples. Statistics were performed in GraphPad Prism. Statistical significance was determined by one-way ANOVA and two-ANOVA followed by pairwise student’s *t*-test with Tukey’s or Sidak’s correction. Two-tailed unpaired student’s *t* test was performed if only two conditions were compared. Data in the text is presented as mean ± s.d. Adjusted P values are reported in the figures.

### Reporting Summary

Further information on research design is available in the Nature Research Reporting Summary linked to this paper.

### Data availability

All data generated and/or analysed during this study are either included in this article (and its Supplementary Information) or are available from the corresponding author on reasonable request. Source Data for Figs. 1, 2 and 4 and Extended Data Fig. 2 and 9 are provided with the article.

## ACKNOWLEDGMENT

The work in Munich and Göttingen was supported by grants from the German Research Foundation (SPP2191, TRR128-2, TRR274, SyNergy Excellence Cluster, EXC2145, Projekt ID390857198, EXC 2067/1-390729940, and STA 1389/5-1), the ERC (Consolidator Grant to M.S.), and the Dr. Miriam and Sheldon G. Adelson Medical Research. The work at the University of Helsinki was supported by the University of Helsinki and by donations of Finnish colleagues to whom we are very grateful. The academy of Finland supported GB, RO and KK (318434), JH and LS (1308613 and 1314119). MJ is supported by The Australian Research Council’s Discovery Early Career Researcher Award (DE190100565). FAM is supported by an Australian National Health and Medical Research Council Senior Research Fellowship (GNT1060075). TT and AT are supported by the European Regional Development Fund (Project No. 2014-2020.4.01.15-0012), by Welcome Trust International Fellowship WT095077MA, by European Research Council grant GLIOGUIDE and Estonian Research Council (grants PRG230 and EAG79, to T.T.). We thank René Müller, Katja Schulz and Uta Scheidt for expert technical assistance.

## AUTHOR CONTRIBUTION

G.B., M.S., A.H. conceived the project. L.C.C., R.O., L.D.P, M.D., J.F., S.K., K.K., M.A., L.S. designed and carried out experiments, A.T., T.T., L.L., O.V., J.H., H.K.K., P.O., M.J., developed and provided tools, L.C.C., R.O., L.D.P., M.D., J.F., S.K., T.K., O.G., C.S., T.S., M.J., F.A.M., S.B., J.H., O.V. analyzed the data or supervised data acquisition. L.C.C., R.O., L.D.P., M.D., J.F., S.K., T.K., O.G. visualized the data, M.S.W., B.M., C.S., H.K.K. provided human samples, G.B. and M.S. wrote the manuscript, G.B. and M.S. supervised the project.

## COMPETING INTEREST

The authors have no competing interests.

## ADDITIONAL INFORMATION

### EXTENDED DATA FIGURES

**Extended Data Fig. 1.**
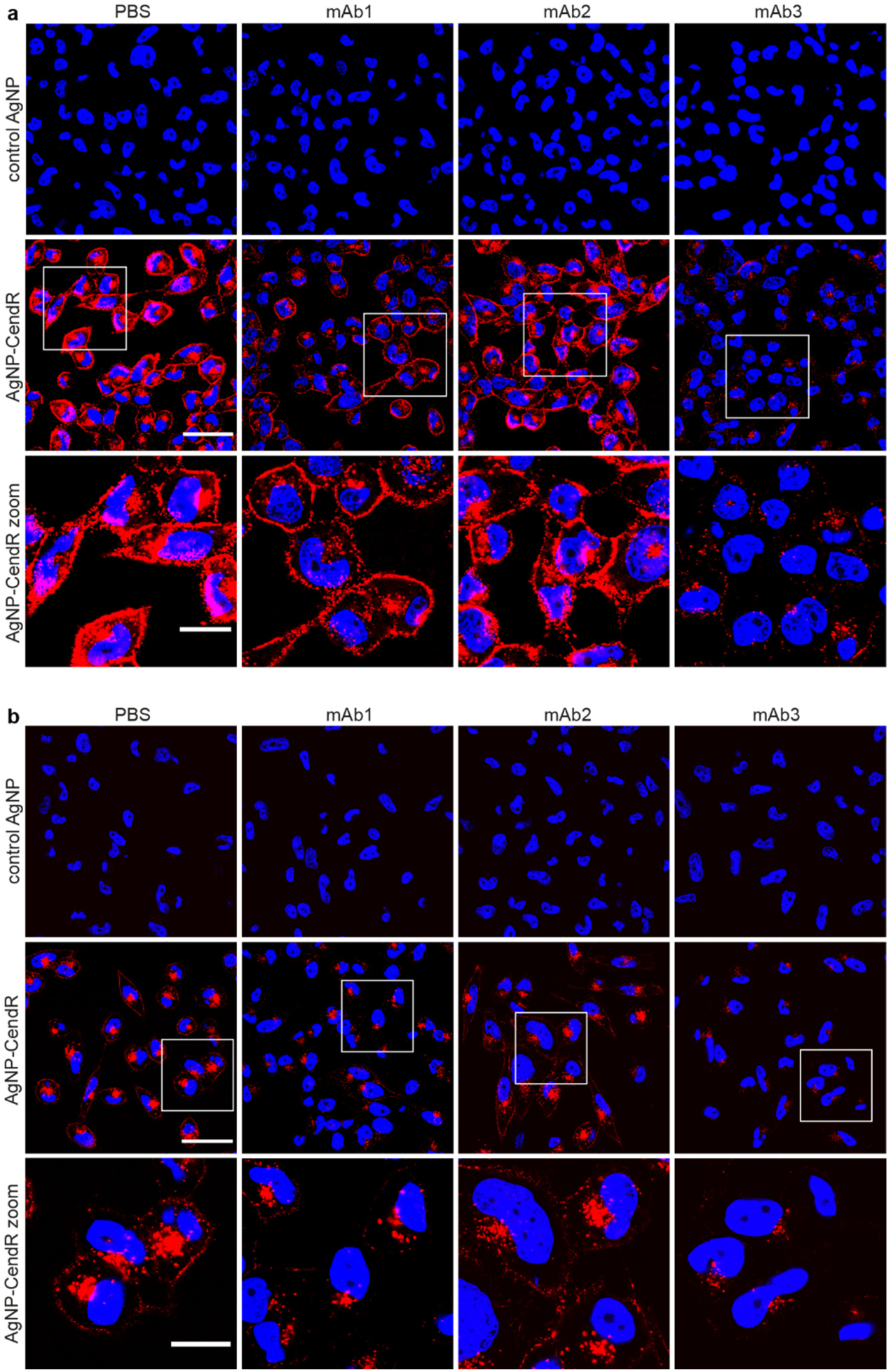
Functional validation of monoclonal blocking antibodies against NRP1. **a**, Representative images of PPC1 cells incubated for 1h with CF555-labeled silver nanoparticles (AgNP) conjugated with biotin (control AgNP, top row) or biotin-X-RPARPAR (AgNP-CendR, bottom row). Cells were pre-treated with a vehicle control (PBS) or the indicated mAb against NRP1 showing the binding and internalizing of the AgNP-CendR nanoparticles. **b**, PPC1 cells treated as in (**a**) were incubated with an etching solution 5 min before fixation to solubilize the non-internalized nanoparticles bound to the external side of the cells. AgNP (red), DAPI (blue). Scale bars, 50 µm and 20 µm (zoom).

**Extended Data Fig. 2.**
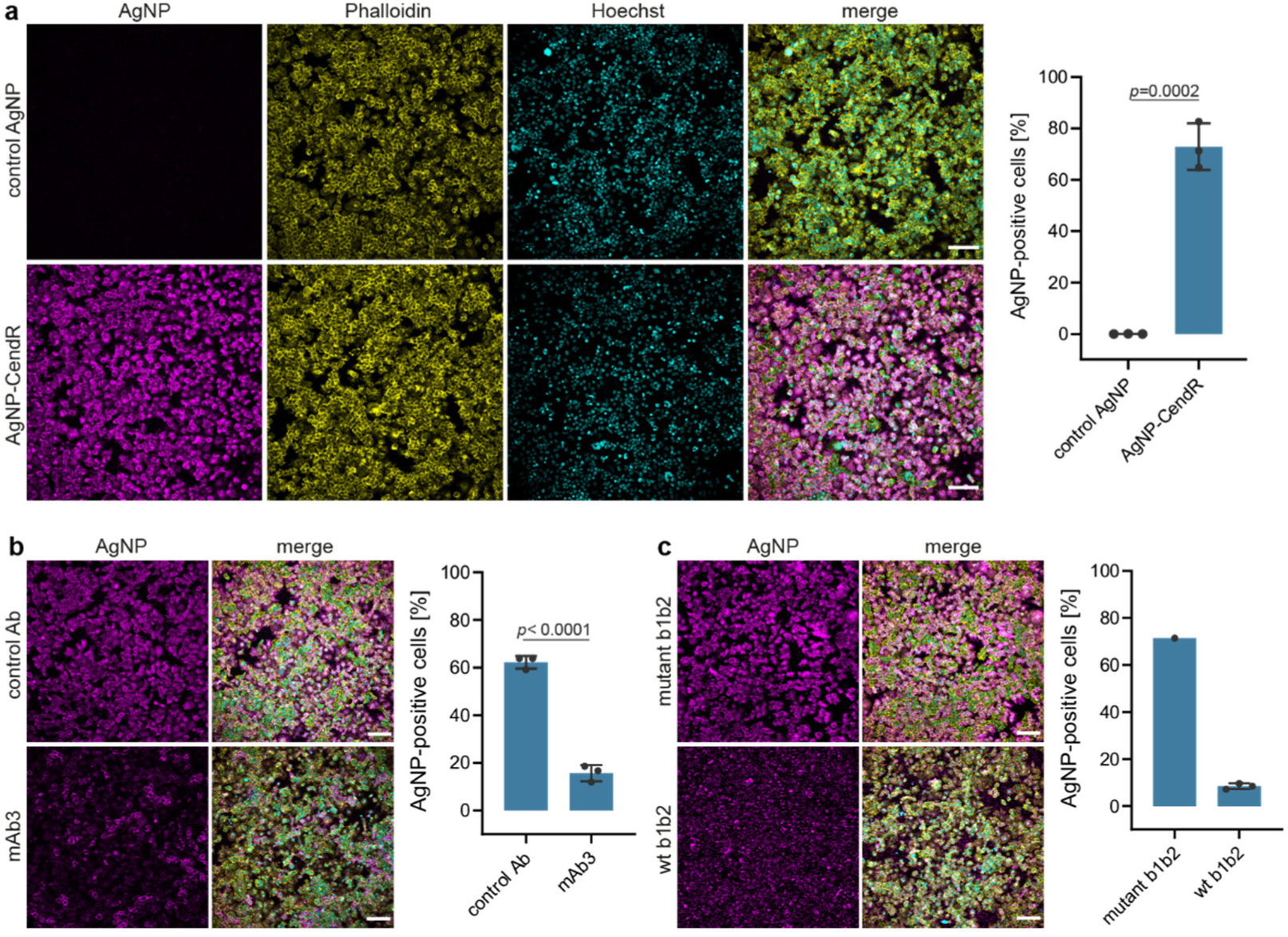
Anti-NRP1 monoclonal antibody (mAb3) or the b1b2 domain (wt b1b2) of NRP1 suppress the NRP1-dependent uptake of labelled silver nanoparticles (AgNPs). **a**, Representative images and quantification showing the uptake of CF647-labeled AgNP conjugated with biotin-X-RPARPAR (AgNP-CendR, lower row) or biotin alone (control AgNP, upper row) after 30 minutes in HEK-293T cells expressing NRP1. AgNPs (magenta) were visible inside the cells, counterstained with Alexa-Fluor488-phalloidin (yellow) and Hoechst (cyan). **b**, Representative images and quantification of NRP1-expressing HEK-293T cells treated for 30 minutes with the blocking mAb3 antibody (lower row) or control Ab (upper row). **c**, Representative images and quantification of NRP1-expressing HEK-293T cells incubated for 30 minutes with AgNPs, together with wild-type b1b2 domain (wt b1b2, lower row) or the mutant b1b2 domain of NRP-1 (mutant b1b2, upper row). *n* = 3 mice. Data are means ± s.d. Two-tailed unpaired Student’s t test. Scale bars, 100 µm.

**Extended Data Fig. 3.**
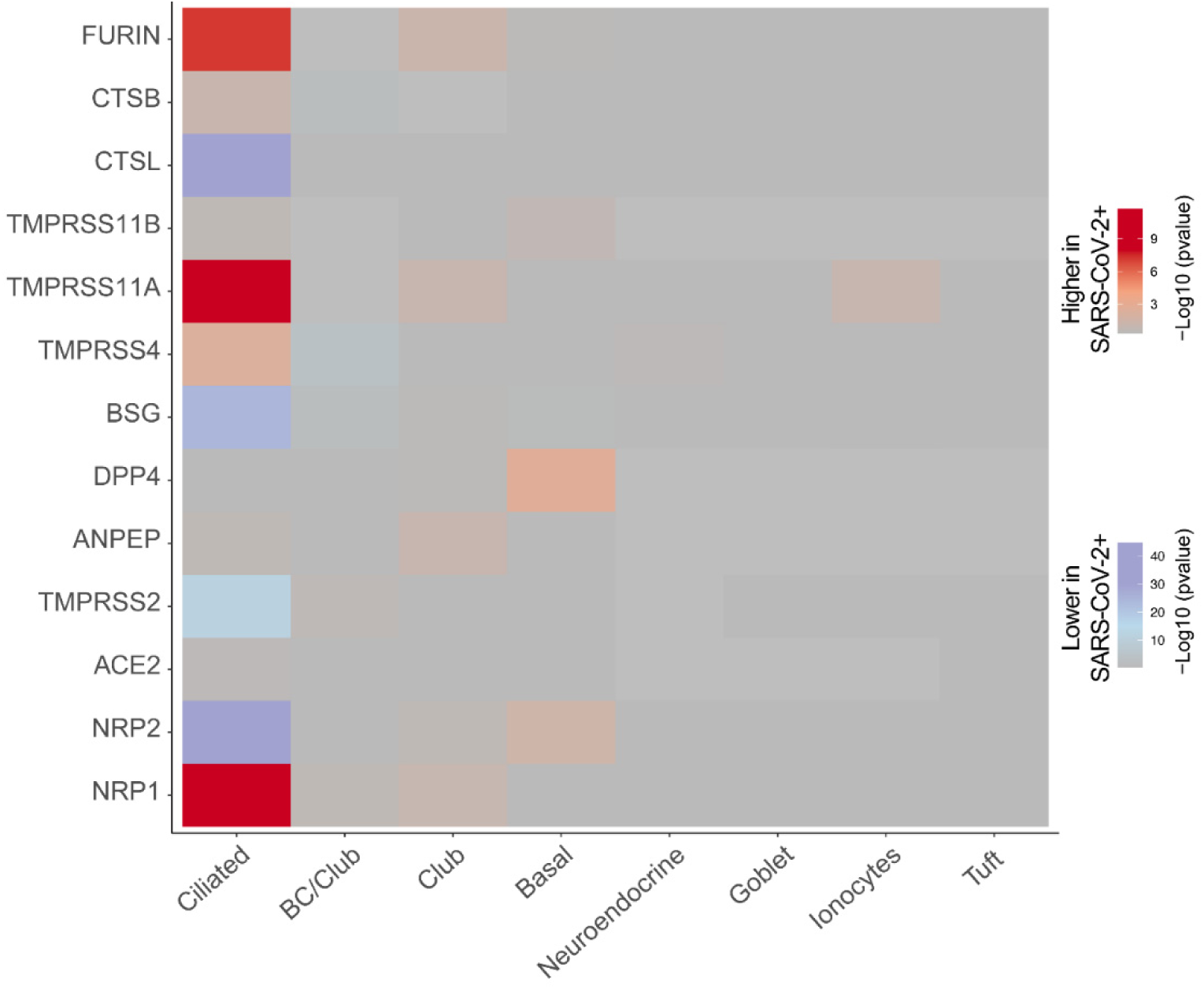
Comparing known SARS-CoV-2 entry determinants across infected versus non-infected bronchial epithelial cells. Navindra *et al*.^16^ infected human bronchial epithelial cells (HBECs) with SARS-CoV-2 and used scRNA-seq to identify ciliated cells as the major target of infection. Among the proposed cell entry and amplification factors only NRP1, Furin and TMPRSS11A are highly enriched in SARS-CoV-2 positive ciliated cells.

**Extended Data Fig. 4.**
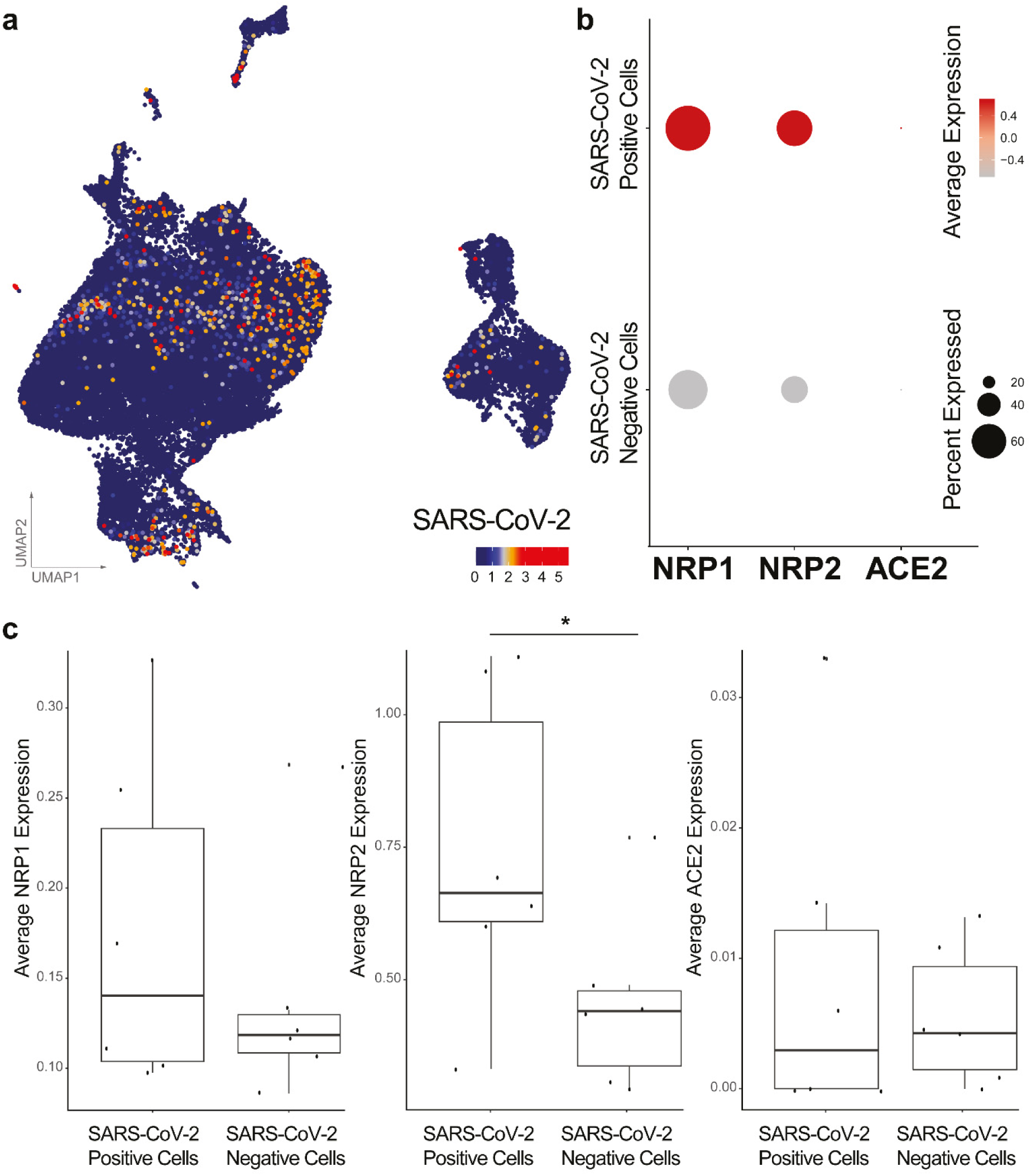
RNA expression of NRP1, NRP2 and ACE2 in SARS-CoV-2 positive cells isolated from patient bronchoalveolar lavage fluids. **a**, UMAP dimensionality reduction plot of all 63,103 cells with the normalized counts of SARS-CoV-2 reads obtained from Liao *et al*.^17^ **b**, Dot plot visualization of the expression of NRP1, NRP2 and ACE2 in BALF cells. Average expression is the scaled value. **c**, Box plots comparing the average expression levels of NRP1, NRP2 and ACE2 between SARS-CoV-2 positive and negative (bystander) cells from severe cases. Each dot represents a patient with a severe case, Bonferroni-corrected p-value for NRP2 = 3,42E-25 (Wilcoxon rank-sum test).

**Extended Data Fig. 5.**
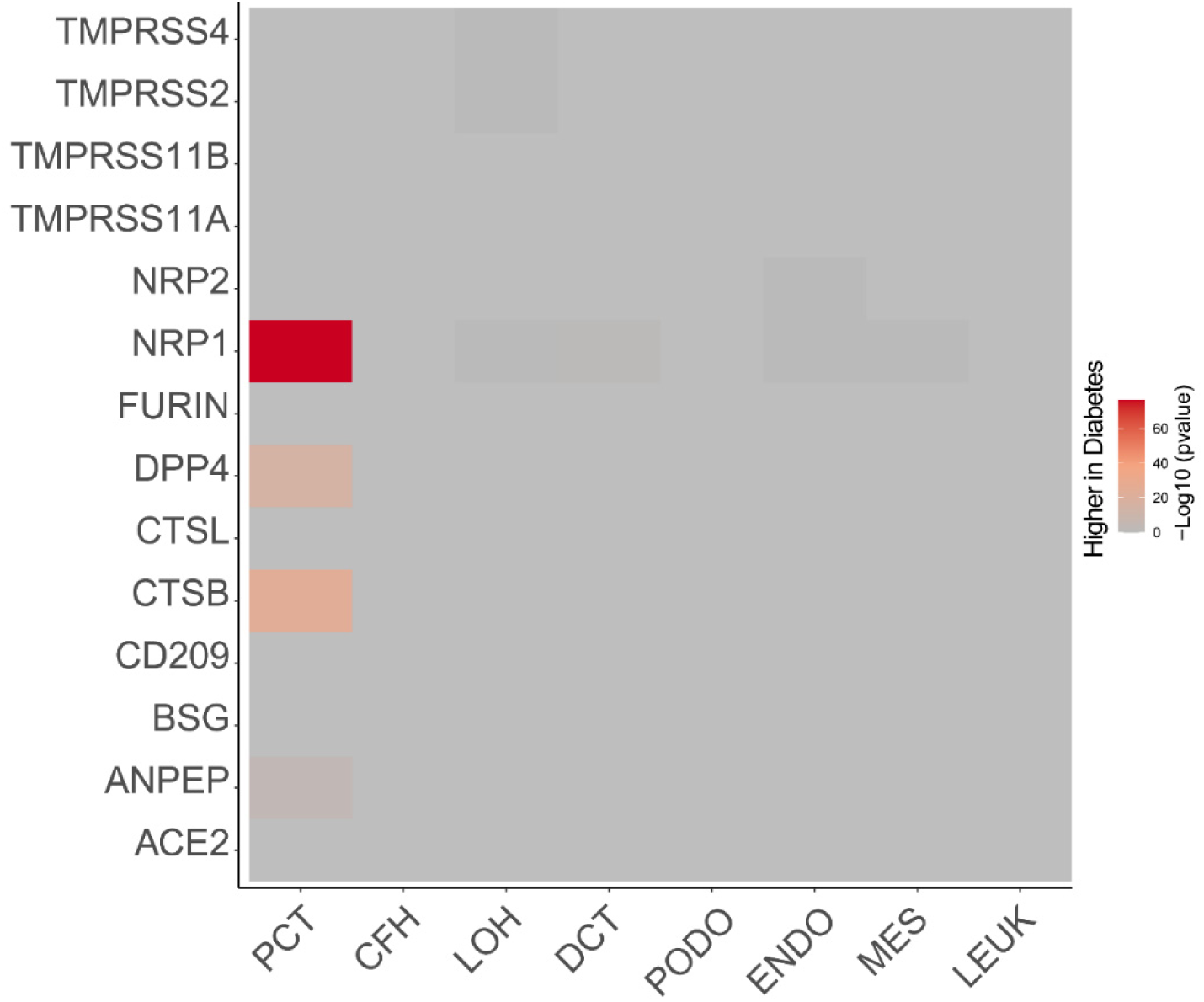
Diabetes induces NRP1 expression in the human kidney. Wilson *et al.*^19^ used single-nucleus RNA sequencing (snRNA-seq) on cryopreserved human diabetic kidney samples to generate 23,980 single-nucleus transcriptomes from 3 control and 3 early diabetic nephropathy samples. Among proposed SARS-CoV-2 cell-entry and amplification factors, only NRP1 is significantly upregulates in PCT (p value= 2,16E-77), proximal convoluted tubule cells. PCT, proximal convoluted tubule; CFH, complement factor H; LOH, loop of Henle; DCT, distal convoluted tubule; CT, connecting tubule; CD, collecting duct; PC, principal cell; IC, intercalated cell; PODO, podocyte; ENDO, endothelium; MES, mesangial cell; LEUK, leukocyte.

**Extended Data Fig. 6.**
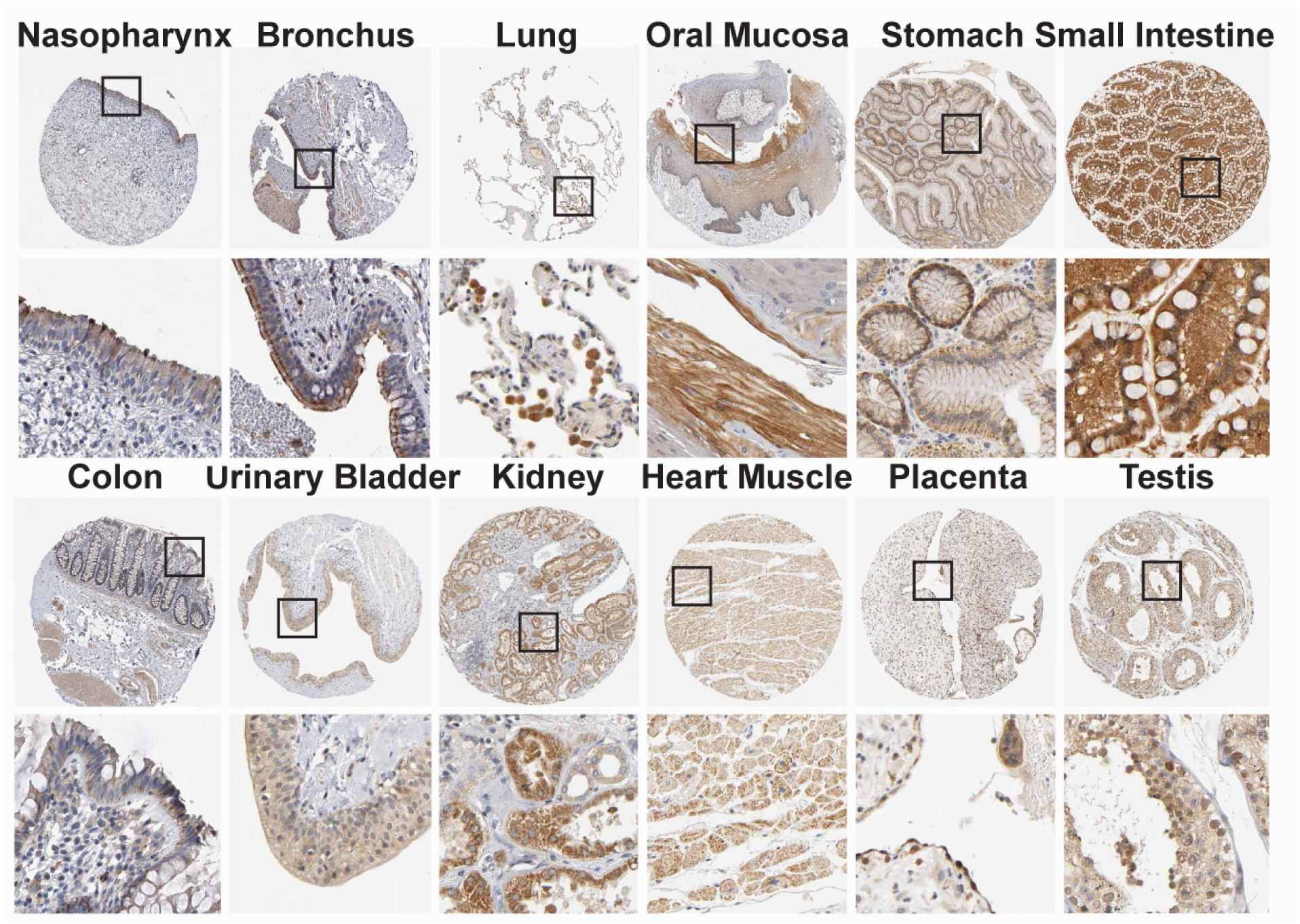
Protein localization of NRP1 in human tissues based on immunohistochemistry. NRP1 immunohistochemistry of 12 human tissues were done using NRP1 antibodies HPA030278 (Atlas Antibodies AB) or CAB004511 (Santa Cruz Biotechnology) (brown), and counterstained with hematoxylin (blue). Particularly high expression of NRP1 was observed in many organs in the epithelial cells facing the external environment. For details, see https://www.proteinatlas.org/ENSG00000099250-NRP1/tissue/primary+data.

**Extended Data Fig. 7.**
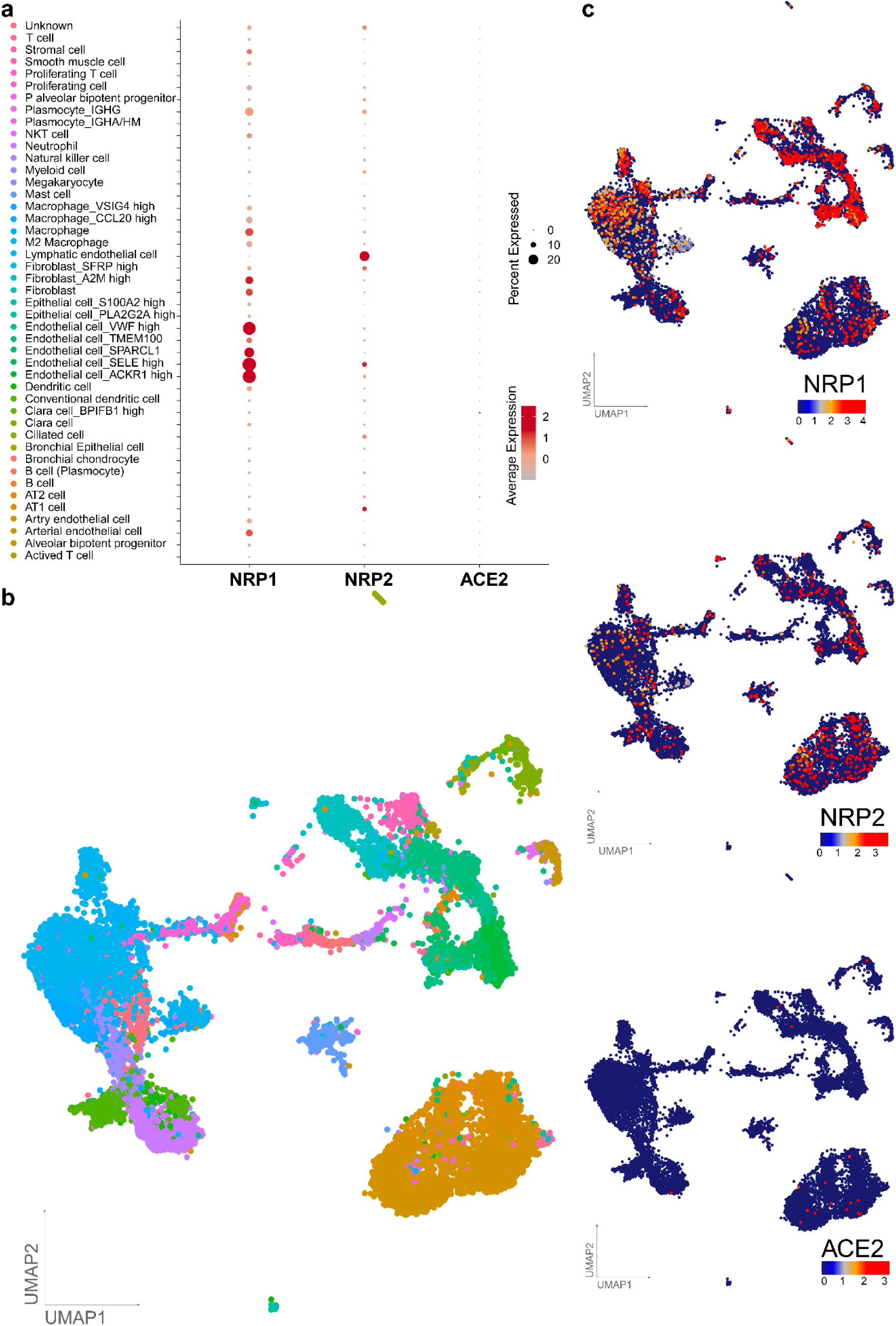
RNA expression of NRP1, NRP2 and ACE2 in adult human lung cells. **a**, Dot plot visualization of the expression of NRP1, NRP2 and ACE2 in lung cell subtypes. The size of the dots indicates the proportion of cells in the respective cell type having greater-than-zero expression of NRP1 (first column), NRP2 (second column) and ACE2 (third column) while the color indicates the mean expression of aforementioned genes. Average expression is the scaled value. **b**, UMAPs are showing the lung cell subtypes and expression levels of NRP1, NRP2 and ACE2. Each dot represents a single cell (cell number n = 17,438). The cell cluster identity is noted on the color key legend and labels based on Han *et al.*^21^.

**Extended Data Fig. 8.**
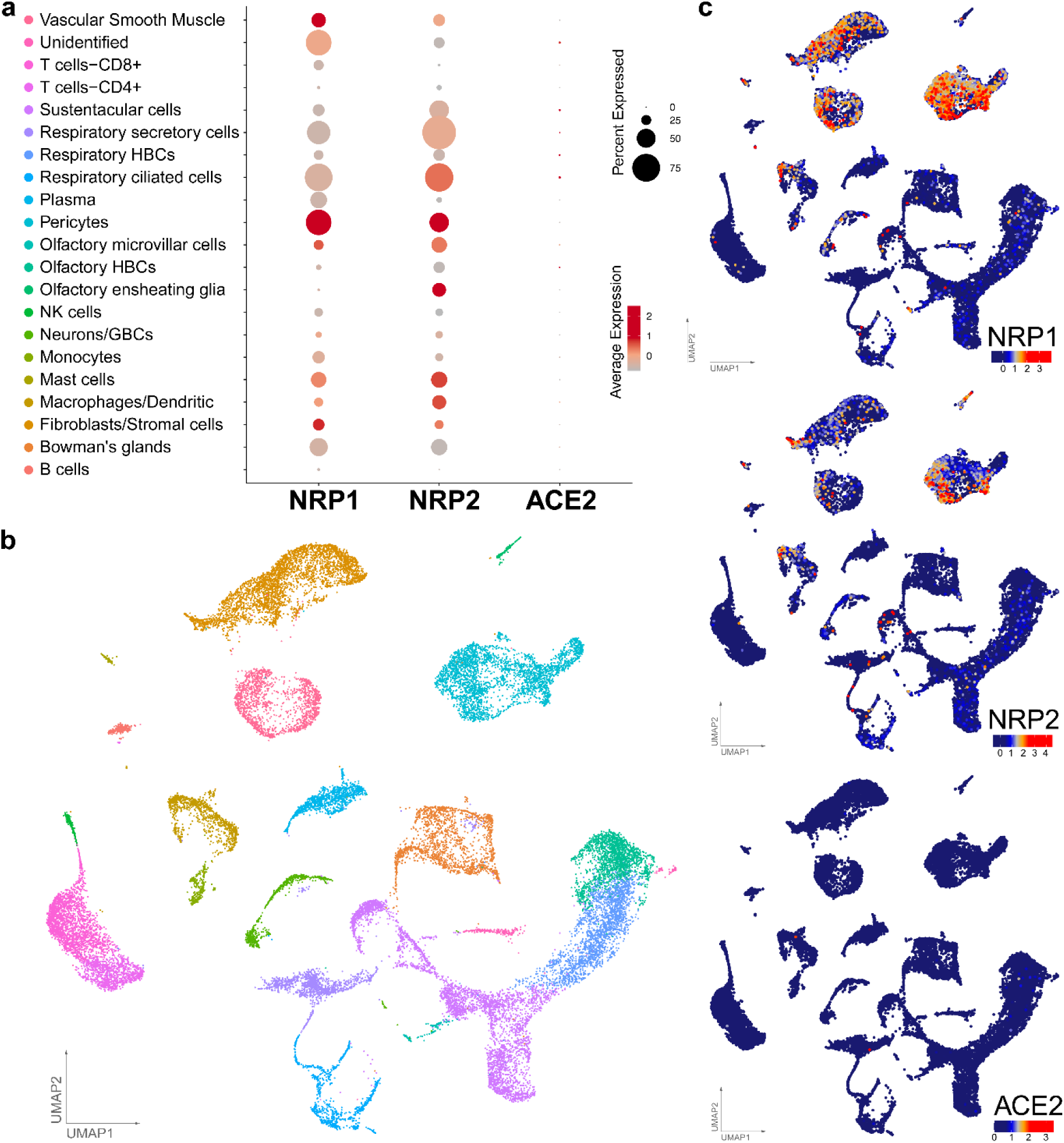
RNA expression of NRP1, NRP2 and ACE2 in the adult human olfactory neuroepithelium. **a**, Dot plot visualization of the expression of NRP1, NRP2 and ACE2 in olfactory neuroepithelia. The size of the dots indicates the proportion of cells in the respective cell type having greater-than-zero expression of NRP1 (first column), NRP2 (second column) and ACE2 (third column) while the color indicates the mean expression of aforementioned genes. Average expression is the scaled value. **b**, UMAPs depicting the olfactory neuroepithelial cell types (cell number n = 28,622) and expression levels of NRP1, NRP2 and ACE2. The cell cluster identity is noted on the color key legend and labels based on Durante *et al.*^22^.

**Extended Data Fig. 9.**
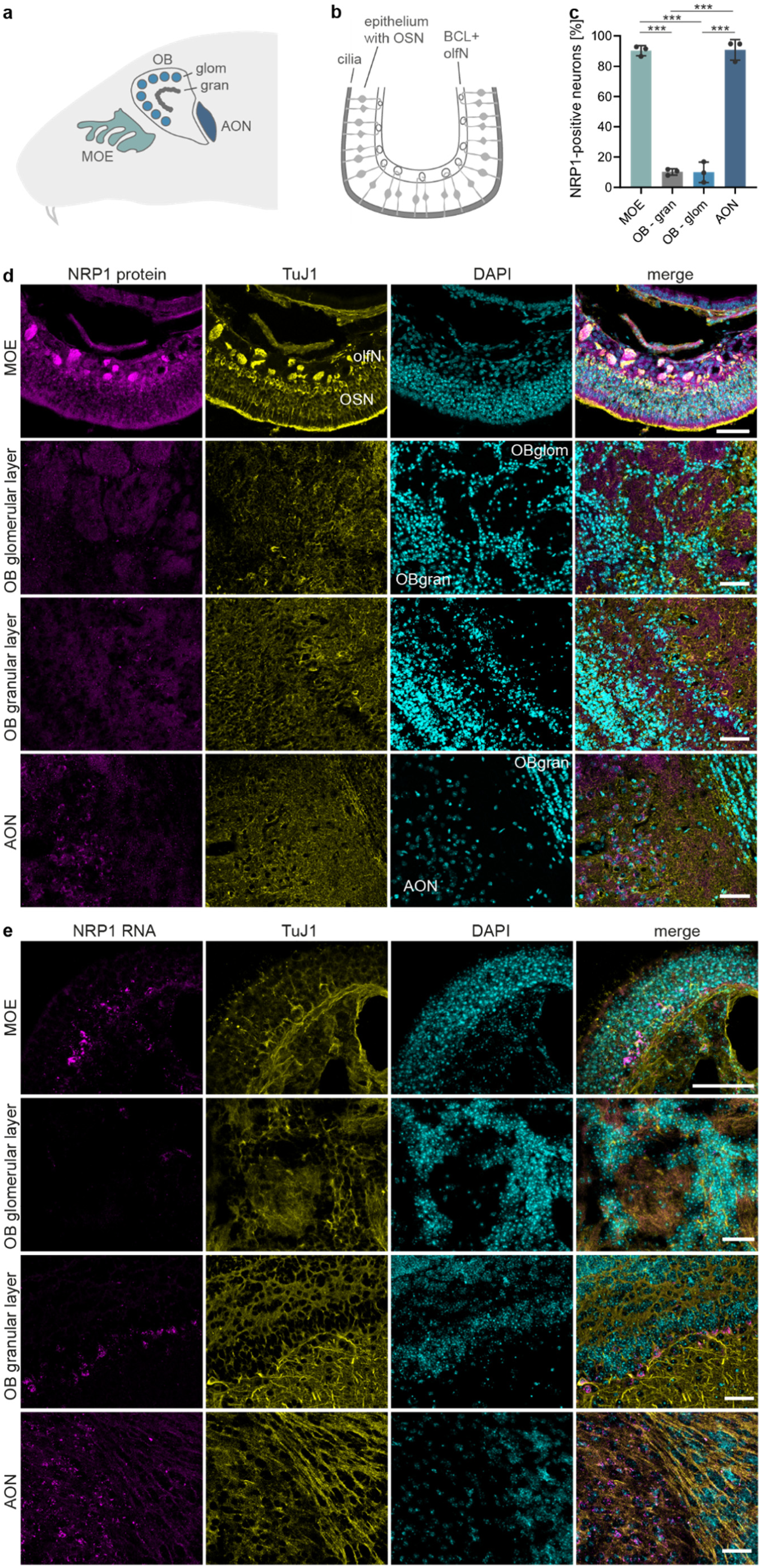
NRP1 is expressed in the mouse main olfactory epithelium (MOE) and the olfactory bulb (OB). **a**, Schematic representation of the mouse head showing the position of the main olfactory epithelium (MOE), the glomerular (OBglom) and granular (OBgran) layers of the olfactory bulb (OB), and the anterior olfactory nucleus (AON). **b**, Schematic representation of the MOE, depicting olfactory sensory neurons (OSN) within the epithelium, their apical cilia, and olfactory nerves (olfN) within the basal cell layer (BCL). **c**, Quantification of NRP1^+^ neurons in the mouse MOE, OB and AON. *n* = 3 mice. Data are means ± s.d. One-way ANOVA with Tukey’s correction for multiple comparisons. **d**,**e**, Confocal images of NRP1 protein (upper panels) and NRP1 RNA (lower panels) in the mouse olfactory system, showing co-localization of NRP1. Pseudocolored images, NRP1 protein and NRP1 RNA (magenta), TuJ1 (yellow), Hoechst (cyan). Scale bars, 50 μm. \*\*\**p* < 0.0001.

**Extended Data Fig. 10.**
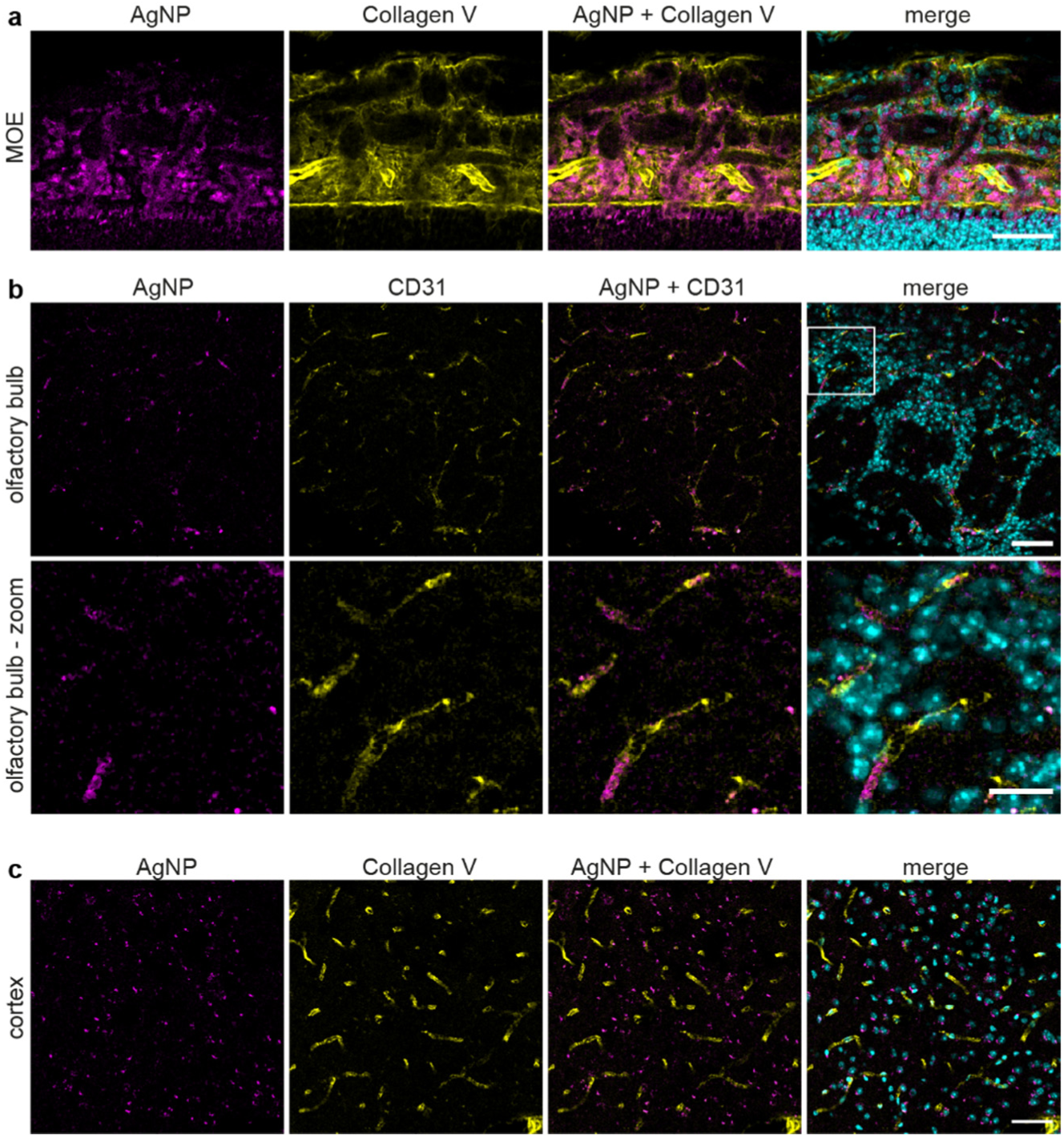
The C-end terminal peptide (CendR) mediates the NRP1-dependent uptake to the blood vessels. **a**, Representative images of main olfactory epithelium (MOE) of animals treated for 6 hours with CF647-labeled AgNPs coated with X-RPARPAR (AgNP-CendR) showing the co-localization between blood vessels (Collagen V, yellow) and the nanoparticles (magenta). **b**, Representative images (upper row) and magnification (lower row) of the olfactory bulb (OB) from AgNP-CendR treated animals showing the co-localization between AgNP-CendR (magenta) and endothelial cells (CD31, yellow). **c**, Representative images of the mouse cortex from AgNP-CendR treated animals showing that the majority of the particles were uptaken by the surrounding cells and can no longer be observed in the blood vessels (collagen V, yellow). Hoechst (cyan). Scale bars: a, 50 μm; b, 50 μm (upper) and 20 μm (zoom), c, 50 μm.

**Extended Data Table. 1.** Patient characteristics of COVID-19 (1 to 6) and control patients (7 to 13).

| ID | Gender | Age | Clinical history | Spike Protein |  | SARS-CoV-2 |
| --- | --- | --- | --- | --- | --- | --- |
|  |  |  |  | OE | OB | PCR |
| 1 | m | 71 | aHT, Parkinson's disease, prostate hyperplasia | + | + | + |
| 2 | m | 80 | aHT, Afib, endocarditis (2014), stage 4 CKD, anemia | + | + | + |
| 3 | f | 72 | - | + | + | + |
| 4 | f | 98 | aHT, Afib, bradycardia | + | + | + |
| 5 | m | 68 | - | - | - | + |
| 6 | m | 68 | aHT, Afib, CHD, MI, s/p BAO with MT, T2DM, COPD, stage 2 CKD, AP, bullous pemphigoid | + | + | + |
| 7 | f | 45 | HCC | - | - | - |
| 8 | f | 78 | aHT, Afib, stage 3 CKD, MI | - | - | - |
| 9 | f | 27 | bM, osteoid osteoma (biopsy) | - | n/a | n/a |
| 10 | f | 68 | pituitary adenoma (biopsy) | - | n/a | - |
| 11 | m | 78 | CHD, Parkinson's disease | n/a | - | n/a |
| 12 | m | 45 | MI | n/a | - | n/a |
| 13 | f | 67 | Lewy-Body dementia, anemia | n/a | - | n/a |

Abbreviations: Afib = atrial fibrillation, aHT = arterial hypertension, AP = acute pancreatitis, BAO = basilar artery occlusion, CHD = chronic heart disease, CKD = chronic kidney disease, COPD = chronic obstructive pulmonary disease, GPA = granulomatosis with polyangiitis, HCC=hepatocellular carcinoma, MI = myocardial infarction, MT = mechanical thrombectomy, Ob = olfactory bulb, OE = olfactory epithelium, s/p = status post, T2DM= type 2 diabetes mellitus.

## Notes

### Competing Interest Statement

The authors have declared no competing interest.

